# Estradiol mediates stress-susceptibility in the male brain

**DOI:** 10.1101/2022.01.09.475485

**Authors:** Polymnia Georgiou, Ta-Chung M. Mou, Liam E. Potter, Xiaoxian An, Panos Zanos, Michael S. Patton, Katherine J. Pultorak, Sarah M. Clark, Vien Ngyuyen, Chris F. Powels, Katalin Prokai-Tatrai, Istvan Merchenthaler, Laszlo Prokai, Margaret M. McCarthy, Brian N. Mathur, Todd D. Gould

**Author notes:** <u>**Correspondence to**</u>: Todd D. Gould, MSTF 936, 685 W. Baltimore Street, University of Maryland, Baltimore, MD, 21201. Current address: Department of Psychology, University of Cyprus, Nicosia, Cyprus (Panos Zanos).

## Abstract

In susceptible populations, stress is a major risk factor for the development of mental disorders, including depression. Estradiol, often considered a female hormone, is distributed in the male brain via aromatization of testosterone. The role of estrogen receptors (ERs) in male stress susceptibility and depression is not well understood. We found that absence of ERβ is associated with susceptibility to stress in male mice and that activity of ERβ-projecting neurons from the basolateral amygdala to nucleus accumbens is reduced in hypogonadal mice subjected to stress, while activation of this circuit reverses stress-induced maladaptive behaviors. We identified that absence of estradiol, but not testosterone *per se*, underlies stress susceptibility and that brain-selective delivery of estradiol prevents the development of depression-related behaviors. Our findings provide evidence for an estrogen-based mechanism underlying stress susceptibility and offer an unexpected therapeutic strategy for treating depression in males.

## Main text

17β-estradiol (E2) is a gonadal steroid hormone with actions in different tissues including the uterus, anterior pituitary, skeletal muscles, as well as the central and peripheral nervous systems. Although E2 is commonly considered as the “female hormone”, it is well distributed in the male brain as a consequence of testosterone’s conversion to E2 by the actions of the aromatase enzyme (*1*–3). Fluctuations in E2 levels are associated with an increased risk for development of mood disorders in women (*4*) and E2 administration enhances reward sensitivity in female rodents (*5*–7), suggesting a possible role of this hormone in reward-deficit associated disorders such as depression. Indeed, E2 administration to women exerts antidepressant effects (*8*–10). Similarly, testosterone exerts antidepressant effects in males (*11*–13); however, the role of E2 in males has received limited attention due to the assumption that the demonstrated antidepressant effects of testosterone in males are through the actions of testosterone itself. We hypothesized that testosterone acts as a precursor and that it is E2 that is responsible for testosterone’s antidepressant effects in males through its action on estrogen receptors (ERs).

### ERβ is involved in the development of maladaptive behaviors in males following exposure to mild stress

We assessed whether lack of ERα or ERβ in conventional knockout mice (ERKO or BERKO, respectively) results in maladaptive stress-sensitive behaviors. Under baseline conditions we found that neither a lack of ERα (Extended Data Fig 1A-E) nor ERβ (Fig Extended Data Fig 1F-J) induce robust behavioral changes in gonadally-intact male mice. We assessed behaviors following subthreshold social defeat stress, which is a single day social stress procedure where experimental mice are introduced to one side of a divided home cage of an aggressor for two minutes, then separated on the opposite side for 15 minutes, and repeated three times. This procedure does not induce any depressive-related behavioral changes in control mice (*14*) (Fig 1A-E), but we found that male mice with a genetic deletion of ERα manifested susceptibility to develop social interaction deficits (Fig 1B), but not anhedonia as tested by preference for the smell of female *versus* male mouse urine (*15*) (Fig 1C). In agreement with this finding, we found that ERKO mice did not manifest social interaction deficits (Extended Data Fig 1K) or sucrose preference deficits (*16*) following inescapable foot shock stress. In contrast, genetic deletion of ERβ increased susceptibility to develop both social interaction deficits (Fig 1D) and anhedonia (Fig 1E) in male mice, suggesting that ERβ might play a greater role in stress susceptibility than ERα. These maladaptive phenotypes were not observed in control ERKO or BERKO mice that were not subjected to stress (Fig 1B- E), indicating that a stress episode, albeit relatively mild, is essential to reveal a susceptible depressive phenotype induced by the lack of ERβ. Social interaction deficits and anhedonia were also induced by foot shock stress in BERKO mice (Extended Data Fig 1L-M), indicating that the ERβ mediated stress susceptibility observed is not dependent on a specific type of stress. In line with these findings, we have previously shown that BERKO, but not ERKO mice, demonstrate decrease in sucrose preference following foot shock stress (*16*), suggesting that the observed effects are not due to the effects of ERβ on chemo-investigation (*17*). In support of this conclusion, non-stressed BERKO male mice demonstrated the expected female urine preference, suggesting that the lack of preference is due to stress and not due to other confounding factors (Fig 1E). While we demonstrated that ERβ contributes to susceptibility in males, we found that lack of ERβ in female mice imparts stress resilience following inescapable foot shock induced social interaction deficits (Extended Data Fig 1N) and previously reported a similar finding regarding escape deficits (*16*), suggesting a sex-dependent differential phenotype and warrants further investigation.

**Fig 1:**
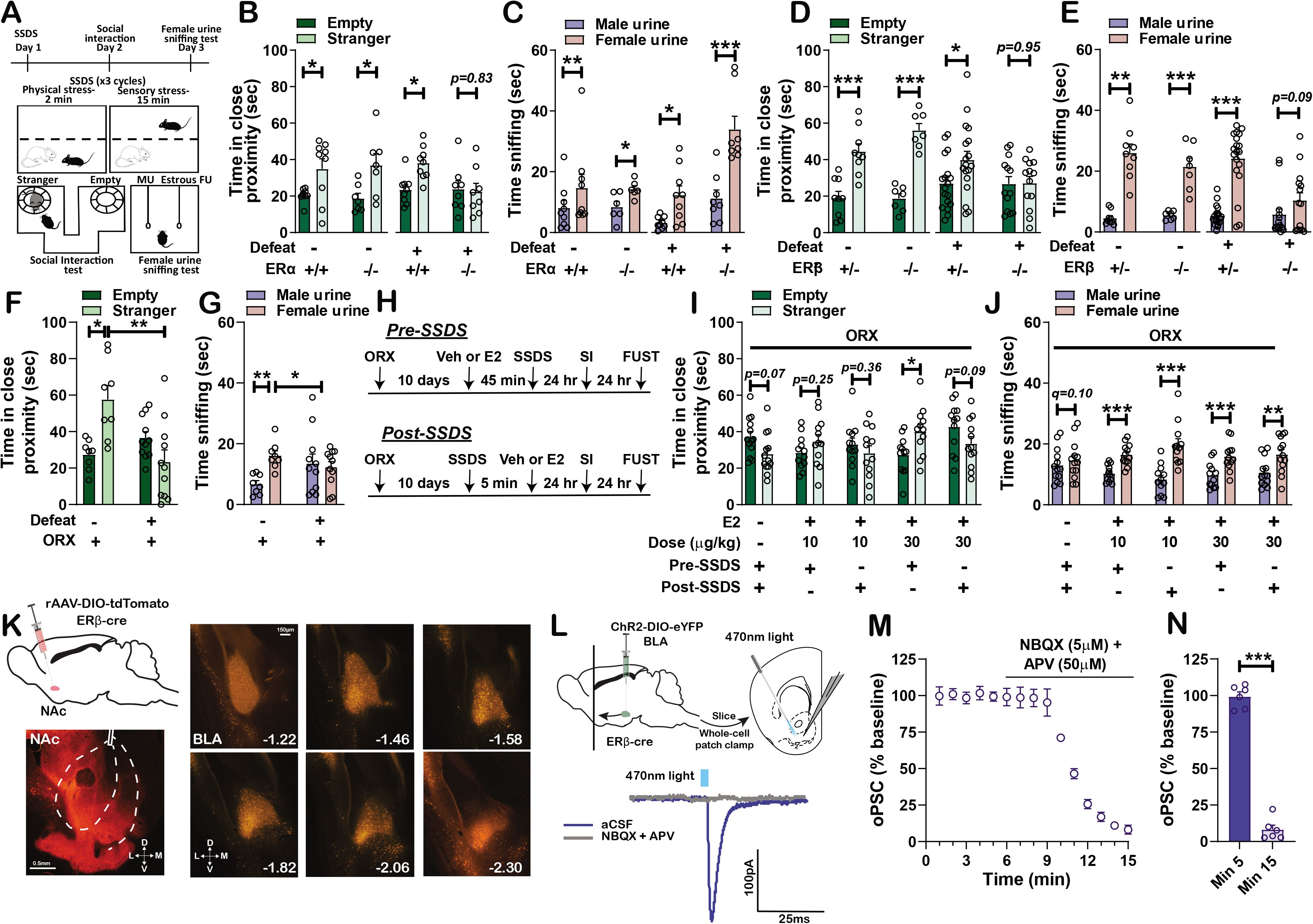
Estrogen receptor beta (ERβ) underlies male stress susceptibility. (A) Timeline and schematics of the subthreshold social defeat stress (SSDS) and behavioral paradigms. Male estrogen receptor alpha knockout mice demonstrated (B) social interaction deficits but not (C) anhedonia following SSDS. Male ERβ knockout mice demonstrated (D) social interaction deficits and (E) anhedonia following SSDS. Orchiectomized (ORX) mice, characterized by the absence of testosterone and consequently estradiol (E2), developed (F) social interaction deficits and (G) anhedonia following SSDS. (H-J) A single s.c. administration of E2 was effective in preventing social interaction deficits and anhedonia when administered 45 min prior to SSDS, while an administration of E2 5 min after SSDS reversed only the observed anhedonia. (K) rAAV-DIO-tdTomato injected in the nucleus accumbens (NAc) of ERβ-cre mice reveals a strong ERβ projection from the basolateral amygdala (BLA). (L-N) Whole-cell patch clamp measurement of optically-induced postsynaptic currents (oPSC) in the NAc of ERβ-cre mice that received an injection of ChR2-DIO-eYFP in the BLA. oPSCs were blocked with bath applied NBQX and APV, indicating that the ERβ-expressing BLA-NAc neurons are glutamatergic. Data shown are the mean ± S.E.M. * *p*<0.05; ** *p*<0.01; \*\*\**p*<0.001. Abbreviations: artificial cerebrospinal fluid, aCSF; (2R)-amino-5-phosphonovaleric acid- APV; basolateral amygdala- BLA; dorsal,D; 17β-Estradiol, E2; estrogen receptor alpha, ERα; estrogen receptor beta, ERβ; female urine- FU; female urine sniffing test, FUST; lateral, L; medial, M; male urine, MU; nucleus accumbens, NAc; 2,3-Dioxo-6-nitro-1,2,3,4-tetrahydrobenzo[*f*]quinoxaline-7-sulfonamide, NBQX; orchiectomy, ORX; optical postsynaptic currents, oPSCs; social interaction, SI; subthreshold social defeat stress, SSDS, ventral, V.

To test whether these behavioral effects were similarly mediated by circulating gonadal hormones, male mice were orchiectomized to remove endogenous testosterone and consequently E2 produced from testosterone and subjected to subthreshold social defeat stress or control conditions. We found that orchiectomized mice manifested the same stress-susceptible phenotype as we observed in the BERKO mice (Fig 1F, G). ORX non-stress controls demonstrated social preference and preference to the female urine (Fig 1F, G), demonstrating that our findings are due to stress and not due to the reduced chemosensory investigation of conspecifics by the loss of hormones as previously described (*17*). Moreover, acute administration of E2 prior to mild stress in orchiectomized male mice (for timeline see Fig 1H) prevented the development of both sociability deficits (Fig 1I) and anhedonia (Fig 1J), though administration of E2 after stress reversed only the anhedonia phenotype (Fig 1J). These data reveal that E2 levels and ERβ mediate susceptibility to sub-threshold social stress in male mice.

### Activation of ERβ expressing neurons in the BLA to NAc is rewarding

Next, we endeavored to identify the neural circuit underlying the above-identified role of ERβ in male stress susceptibility. There is ample evidence implicating the nucleus accumbens (NAc) in reward responses and stress resilience/susceptibility (*18*–22). Thus, we investigated whether there are strong ERβ-expressing projections to the NAc by injecting a cre-sensitive retrograde adeno-associated virus into the NAc of mice expressing cre from the endogenous ERβ promoter (ERβ-icre) (see Extended Data Fig 2 for confirmation of expected cre expression). We identified neurons strongly expressing ERβ that project from the basolateral amygdala (BLA) to the NAc (Fig 1K), which was confirmed by injecting a retrograde conjugate cholera toxin B subunit neuronal tract tracer in the NAc of wild-type mice followed by assessing *Esr2* expression co-labelling in the BLA (Extended Data Fig 3A,B) demonstrating that ~70% of the tracer positive cells also express *Esr2*. *Esr2* expression appears to be enriched specifically in the BLA to NAc projection compared with the whole BLA as *Esr2* is expressed at ~10% of the total BLA cells (DAPI labelled) compared with ~26% expression in the BLA to NAc projecting cells (Extended Data Fig 3A,C). Although the known projections from the BLA to NAc are characterized as glutamatergic (*23*–25), ERβ expressing neurons in other amygdala subregions are reported to be exclusively GABAergic (*26*). We demonstrated using whole-cell electrophysiology recordings that the ERβ-expressing BLA to NAc neurons are glutamatergic (Fig 1L-N). Our finding is in line with recent evidence from RNAseq data showing that *Esr2* is expressed in excitatory BLA neurons (*27*). To test whether this projection is involved in stress-susceptibility, mice were orchiectomized, treated acutely with either E2 or vehicle (as in Fig 1 H-J), and underwent subthreshold social defeats followed by assessment for activation of the BLA-to-NAc projection using c-Fos immunochemistry. Prior to E2 administration, mice received an injection of Cre-sensitive rAAV administered to the NAc, which is retrogradely transported to the BLA. Our results demonstrate that following a brief stress exposure, ERβ-expressing BLA-to-NAc neurons are activated at a significantly higher degree in mice that received E2 (Fig 2A-C). Importantly, we did not observe activation of any of the other strong ERβ-projections to the NAc in response to E2, in particular medial prefrontal, cingulate, or insular cortex, suggesting that our effects are specific to the BLA to NAc projections (Extended Data Fig 4A-C).

**Fig 2:**
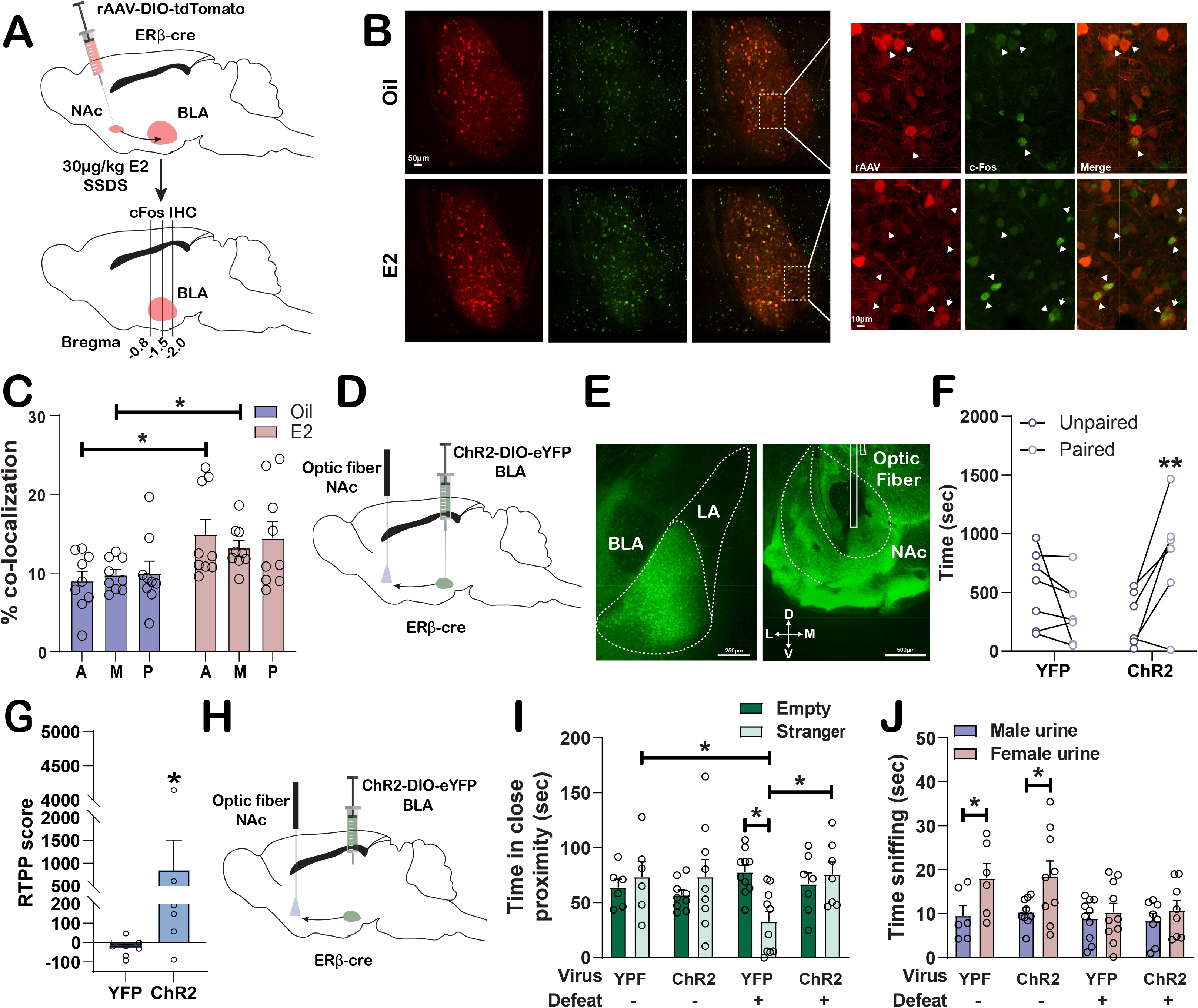
Estrogen receptor beta (ERβ)- expressing basolateral amygdala (BLA) neurons projecting to the nucleus accumbens (NAc) are involved in stress-induced social deficits. (A) rAAV-DIO-tdTomato was injected in the NAc of male ERβ-cre mice and retrogradely transported to cell bodies localized in the BLA. Orchiectomized (ORX) mice received estradiol (E2; s.c.) 45 min prior to subthreshold social defeat stress (SSDS). (B) Representative immunofluorescence images from the BLA showing the co-localization of c-Fos and tdTomato in mice who received either E2 or vehicle (sesame oil) administration. (C) Increased activation of ERβ-expressing BLA to NAc cells in the anterior (A), medial (M), and posterior (P) BLA was observed in mice that received E2 prior to SSDS. (D) Schematic of the ChR2-DIO-eYFP injection in the BLA, optic fiber implantation in the NAc, and (E) representative images from the injection and fiber implantation site. (F-G) Blue light stimulation (470 nm; 20 Hz) of ERβ-expressing BLA to NAc terminals induced a real-time place preference, shown by the increased time spent in the stimulation-paired compartment. (H) Schematic of the ChR2-DIO-eYFP injection in the BLA and optic fiber implantation in the NAc. Blue light stimulation (470 nm; 4Hz) of ERβ-expressing BLA to NAc terminals for 45 min prior to stress in ORX mice prevented (I) social interaction deficits but not (J) anhedonia. Abbreviations: Anterior, A; basolateral amygdala, BLA; channelrhodopsin-2, ChR2; 17β-Estradiol, E2; estrogen receptor beta, ERβ, immunohistochemistry, IHC; middle, M; nucleus accumbens, NAc; orchiectomy, ORX; posterior, P; subthreshold social defeat stress, SSDS.

We then investigated the possible involvement of the ERβ-neuronal projection from the BLA to NAc in reward-related behaviors by assessing place preference induced by optogenetic activation of this circuit. Optogenetic activation of the ERβ-expressing BLA-to-NAc terminals when animals entered one of two compartments induced robust real-time place preference in male (Fig 2D-G) but not female (Extended Data Fig 5) ERβ-cre mice that received ChR2-DIO in the BLA and blue light stimulation to the NAc terminals when in the assigned compartment, while control, YFP-injected control mice did not develop real-time place preference (Fig 2D-G). To investigate whether manipulation of this projection might underlie our observed stress susceptibility behavioral effects, mice received an injection of ChR2 or YFP in the BLA (Fig 2H) and were orchiectomized prior to subthreshold social defeats. Light activation of the ERβ-expressing BLA-to-NAc terminals prevented stress-induced social avoidance in orchiectomized mice, while no effect was observed in non-stressed controls, or in the FUST (Fig 2I, J).

We thus hypothesized that in the absence of E2 and following stress exposure, terminal activity of ERβ neurons projecting from BLA to NAc would be decreased during social interaction. To test this hypothesis, we performed *in vivo* fiber photometry measurements in awake-behaving orchiectomized mice following subthreshold social defeat stress during the social interaction test. This was accomplished by injecting Cre-dependent virus expressing GCaMP6s into the BLA of ERβ-Cre mice and recording calcium transients in the NAc terminals thus providing specificity in recording from only the ERβ- projecting BLA neurons (Fig 3 A,B). We determined that during interaction with a different male mouse, axon terminal calcium activity is decreased in orchiectomized mice which received prior stress, compared to sham controls (Fig 3C-I). Interestingly, our data demonstrate that ERβ BLA to NAc neurons remain partially responsive to the stranger mouse, albeit at lesser extent compared with intact mice (Fig 3G), suggesting that these neurons did not lose their ability to respond but rather that the reward value of the stranger mouse interaction decreased. In gonadally intact mice we observed an increase in calcium transients prior to initiation of social interaction, suggesting that this neuronal response might be due to the anticipation of the rewarding stimulus; however, this response was absent in orchiectomized mice (Fig 3E, F, I). These circuit-specific changes were concomitant with social avoidance in orchiectomized mice (Fig 3J, K). We also found that social interaction scores were positively correlated with the calcium signals evoked during social interactions (Extended Data Fig 6A-C). Altogether, these data suggest that the ERβ projection from BLA to NAc has a specific role in E2-related stress susceptibility and development of social impairments.

**Fig 3:**
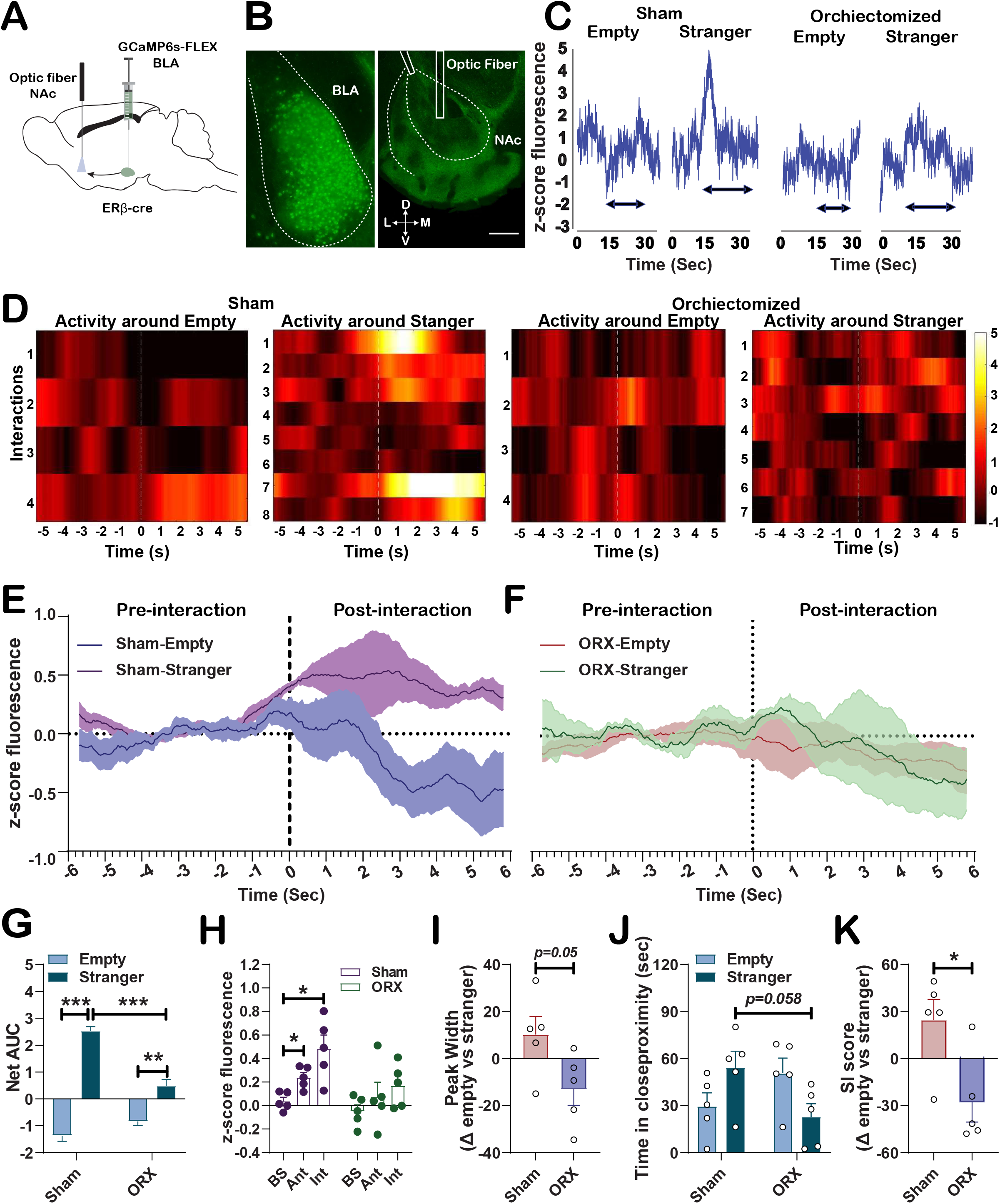
Estrogen receptor beta (ERβ)- expressing basolateral amygdala (BLA) neurons projecting to the nucleus accumbens (NAc) decreased terminal activity is involved in stress-induced social deficits. (A) Schematic of the GCaMP6s-FLEX injection in the BLA, optic fiber implantation in the NAc, and (B) representative images from the injection and implantation sites. (C) Representative traces from the fiber photometry recordings of ERβ-expressing BLA to NAc terminals in which each horizontal arrow represents an interaction bout. (D) Heatmap representation of the changes in z-score fluorescence when gonadally intact and ORX mice were interacting with a stranger or an empty cage. (E-G) Increased calcium transients were observed in ERβ-expressing BLA to NAc terminals while gonadally-intact mice were interacting with the stranger mouse. In contrast, no difference was observed in calcium activity during social interaction in ORX mice. (H) Increased calcium activity was also observed in anticipation of social interaction only in gonadally intact mice. (I) Decreased in peak width calcium activity was observed during social interaction in ORX mice (J-K) Calcium activity in gonadally-intact mice during interaction with the stranger mouse was associated with higher interaction preference compared with ORX mice. Data shown are the mean ± S.E.M. * *p*<0.05; ** *p*<0.01; \*\*\**p*<0.001. Abbreviations: anticipation, Ant; basolateral amygdala, BLA; baseline, BS; estrogen receptor beta, ERβ, interaction, Int; nucleus accumbens, NAc; orchiectomy, ORX; subthreshold social defeat stress, SSDS.

### Absence of estradiol induces stress susceptibility independent of testosterone in males

The above findings have clear relevance to human depression occurring as a consequence of stressful life events. We sought to determine the specific role of chronic circulating hormones in stress susceptibility. First, we demonstrated that orchiectomy on its own, i.e. elimination of gonadal hormones, did not induce any maladaptive behaviors (Extended Data Fig 7A-E), suggesting that similar to humans a second factor like stress is needed to reveal a maladaptive phenotype. Our earlier experiments revealed that, similar to BERKO mice, subthreshold social defeat stress performed in orchiectomized WT mice induced deficits in social interaction (Fig 1F) and female urine sniffing behaviors (Fig 1G). Furthermore, following this mild stress, WT orchiectomized mice manifested deficits in non-social behaviors such as anxiety (Extended Data Fig 7G-I) and short-term memory (Extended Data Fig 7J,K), but no deficits in preference for sucrose solution or struggling time during a forced swimming exposure (Extended Data Fig 7G,M; timeline Extended Data Fig 7F). We then assessed whether testosterone replacement would reverse stress-induced social interaction and hedonic deficits. Mice underwent orchiectomy or sham surgeries and were implanted with either testosterone-filled or empty/control silastic tube implants. Following 10 days of recovery and treatment, mice underwent subthreshold social defeat stress followed by social interaction and anhedonia tests (for timeline see Fig 4A). Testosterone replacement reversed the observed maladaptive behaviors in orchiectomized mice (Fig 4B-D), while no effect of testosterone was observed in gonadally intact or non-stressed mice regardless of gonadal status (Fig 4B-D). Although testosterone replacement therapy is effective for the treatment of refractory-depression in some men (*11*–13), potentially serious life-threatening side effects exist, such as polycythemia and cardiac dysfunction (*28*–30). Thus, potential options that lack testosterone’s side effects and demonstrate specificity of testosterone’s CNS action in mediating stress susceptibility are desirable.

**Fig 4:**
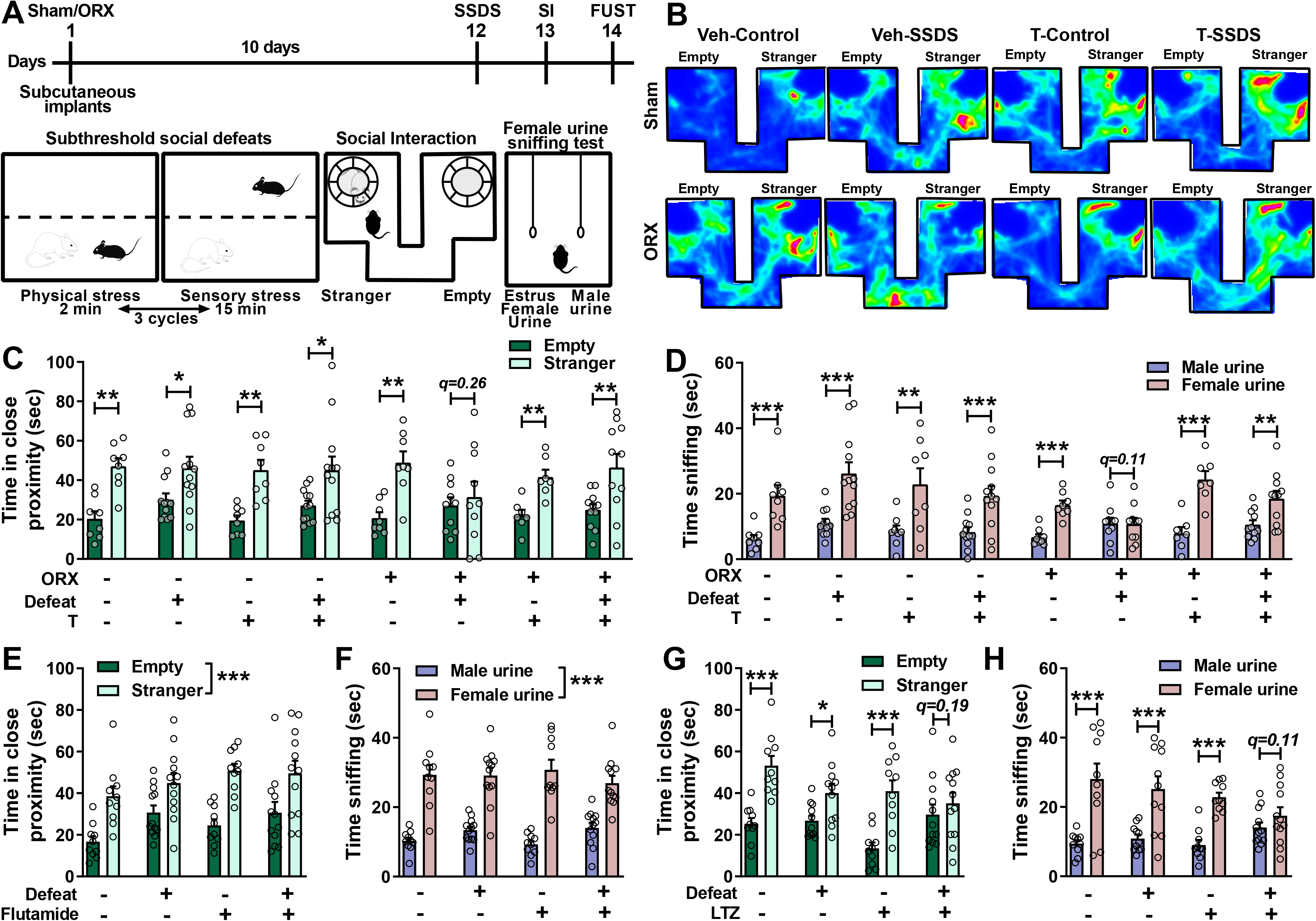
Testosterone does not directly mediate stress susceptibility in male mice. (A) Timeline, and the schematic of the stress and behavioral paradigms for which WT male C57Bl/6J were assessed. (B) Representative heatmap traces during the social interaction test. Chronic administration of testosterone reversed the orchiectomy and SSDS-induced (C) social interaction deficits and (D) anhedonia. In intact mice, blockade of the androgen receptor with chronic flutamide did not induce any deficits in (E) social interaction and (F) anhedonia. Chronic administration of the aromatase inhibitor letrozole in minipumps to gonadally intact mice induced SSDS susceptibility as shown with (G) social interaction deficits and (H) anhedonia. Data shown are the mean ± S.E.M. * *p*<0.05; ** *p*<0.01; \*\*\**p*<0.001. Abbreviations: female urine sniffing test, FUST; letrozole, LTZ; orchiectomy, ORX; social interaction, SI; subthreshold social defeat stress, SSDS; testosterone, T; veh, vehicle

We next demonstrated that blockade of the androgen receptor, which is the only site of testosterone action, with flutamide in intact mice induced neither social deficits (Fig 4E) nor anhedonia (Fig 4F) following stress, bringing into question the direct involvement of testosterone *per se*. Consistent with an E2-specific mechanism of testosterone action, we found that treatment with letrozole, which blocks the aromatization of testosterone to E2 (*31*), induced a stress-susceptible phenotype in intact male mice, as shown by deficits in social interaction (Fig 4G) and anhedonia (Fig 4H). These endophenotypes are consistent with what we observed with hypogonadism, suggesting a direct involvement of E2 in regulating the development of depressive-related behaviors following mild stress in males. In agreement with this conclusion, and our earlier acute administration data (Fig 1H, I), we demonstrated that chronic treatment with E2 through the use of silastic tube implants, reversed the stress-induced social avoidance (Fig 5A) and anhedonia (Fig 5B) in hypogonadal male mice. Moreover, our acute administration data (Fig 1H-J) suggests that E2 can act both as a prophylactic therapy, and, in part, as an antidepressant treatment following the development of the disease. Effectiveness of steroid administration and blockade was confirmed by changes in body, testis and seminal vesicle weights (Extended Data Fig 8A-J, 9A-B). Importantly, hormonal and other manipulations did not affect the aggressiveness of CD1 mice during social defeat bouts as demonstrated by the similar number of attacks (Extended Data Fig 10).

**Fig 5:**
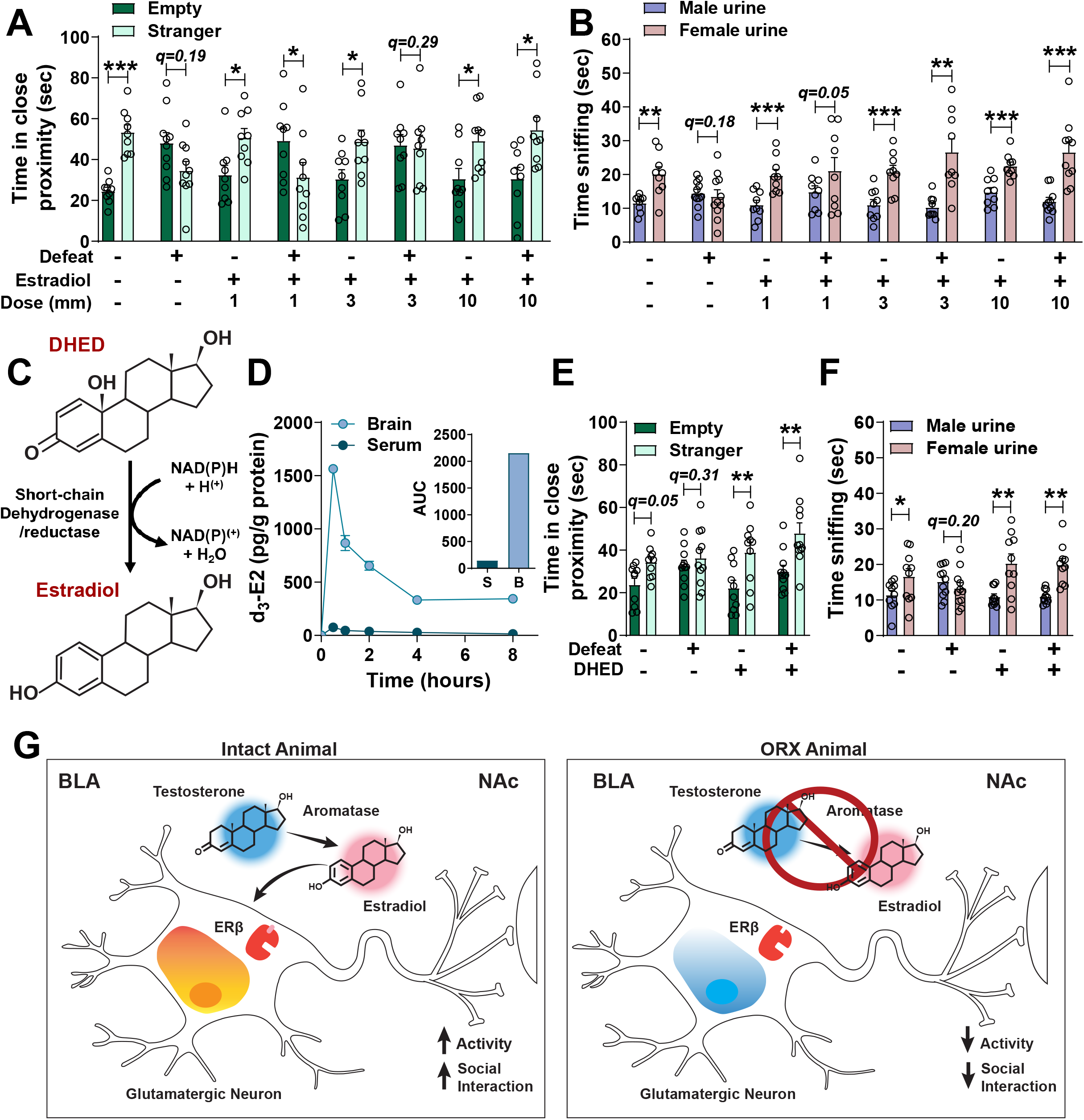
Estradiol (E2) mediates stress susceptibility in male mice. Dose-response following chronic administration of estradiol (E2) reversed the orchiectomy (ORX) and SSDS-induced (C) social interaction deficits and (D) anhedonia. (C) Conversion of a brain selective E2 prodrug DHED to E2. (D) Intact C57Bl/6J male mice that received d3-DHED revealed that DHED derived E2 is found enriched in the brain but only in limited quantities in serum. Chronic administration of DHED with minipumps prevented the SSDS/orchiectomy- induced (E) social interaction deficits and (F) anhedonia. (G) Schematic representation and summary of the suggested mechanism for stress susceptibility in male mice. Data shown are the mean ± S.E.M. * *p*<0.05; ** *p*<0.01; \*\*\**p*<0.001. basolateral amygdala, BLA; (17β)-10,17-Dihydroxy-estra-1,4-dien-3-one, DHED; 17β-Estradiol, E2; estrogen receptor beta, ERβ; orchiectomy, ORX; nucleus accumbens, NAc.

Although our findings support a critical role for E2, and not testosterone itself, as the biologically active hormone mediating stress susceptibility in hypogonadal male mice, E2 replacement therapy cannot be considered as a viable treatment in human male populations due to its peripheral side effects, including gynecomastia and erectile dysfunction (*32*–35). However, isolating the effects of E2 to the central nervous system would provide a viable therapy. It was previously shown that 10β,17β-dihydroxyestra-1,4-dien-3-one (DHED) acts as a prodrug (Fig 5C), that delivers E2 selectively in the brain via conversion of the prodrug by a NADPH-dependent reductase and hence avoiding E2’s peripheral effects (*36*). We performed confirmatory pharmacokinetic analysis in the brain and serum of male mice following peripheral administration of labelled (*d3*)-DHED, where we observed that E2 derived from the labelled DHED is localized to the brain (Fig 5D). Chronic treatment with DHED, in contrast with E2 treatment (Extended Data Fig 9A,B), did not increase body weight and seminal vesicle weight compared with vehicle-treated mice (Extended Data Fig 9C,D) further supporting that DHED lacks E2 peripheral effects. More importantly, we also found that chronic treatment with DHED reversed the social interaction and hedonic deficits induced by the combination of stress and hypogonadism (Fig 5E,F). While E2 administration is not a viable option for the treatment of male depression due to the peripheral effects, our findings provide evidence for an exciting novel approach and suggest viability of future treatments via brain-selective E2 delivery.

Overall, we demonstrated that E2 has a causal relationship and plays a pivotal role in male stress-susceptibility in an animal model relevant to depression. We found that decreased E2 but not testosterone levels *per se* mediates susceptibility to stress in hypogonadal males, through its action on the ERβ. We further demonstrated that ERβ is highly expressed in neurons projecting from the BLA to NAc and that neuronal calcium activity in the NAc terminals of ERβ BLA projection neurons is decreased during social interaction after stress in the absence of E2, and that such changes are correlated with social interaction scores. We identified that stimulation of this circuit rescues social avoidance observed following the combination of stress and hypogonadism (i.e. lack of E2), demonstrating that activity of this circuit is causally related with stress susceptibility (Fig 5G). Our findings provide translationally relevant evidence for the effectiveness of targeting ERs, and in particular ERβ, in the male brain as an effective antidepressant approach. In addition, our studies have elucidated a circuit likely involved in the antidepressant actions of testosterone in hypogonadal males. Although we cannot preclude a role of ERα in stress susceptibility, our findings suggest that genetic deletion of ERβ exerts a stronger stress susceptibility phenotype (Fig 1B-E, Extended Data Fig 1K-M, *16*). In contrast to a prior report demonstrating a role of accumbal ERα in the NAc (*37*), our manipulations focused on the ERβ-expressing cells in BLA to NAc circuit. Earlier experiments by Lorsch et al revealed that ERα does not act as a pro-resilient factor in the PFC, thus suggesting a region-specific role of ERα in stress resilience (37). It is likely ERα exerts its effects by changing local NAc activity, rather than controlling NAc excitability through other projecting regions, such as the PFC or BLA. Altogether, we provide novel evidence for the development of new potential paradigm-shifting and specific therapeutic approaches for the effective treatment of male depression.

## Acknowledgements

We thank by Dr. CheMyong Ko (University of Illinois, Urbana-Champaign) for providing us the ERβ-icre mice to form the breeding colony.

## Author contributions

PG and TDG were responsible for the conceptualization of the project overall experimental design. LEP and PG performed and analyzed the fiber photometry experiments. TMM and KJP performed and analyzed the RNAscope experiments. TMM and SMC performed and analyzed the immunofluorescence experiment. PG and PZ performed the behavioral experiments. CFP performed perfusions and tissue collection. XA sectioned brains for confirmation of viral expression and cannula placement and analyzed PFC, CgCx and InsCx to NAc projections. MSP and BNM performed and analyzed the electrophysiology experiment. VN, LP and KPT synthesized DHED and conducted bioanalytical quantitation of E2 derived DHED. IM and MMC contributed significantly to the design of the hormone manipulation experiments. PG conducted the experiments and their analysis unless otherwise noted. PG and TDG outlined and wrote the paper, which was reviewed by all authors.

## Data and material availability

The authors declare that all data supporting the findings of this study are available within the paper. Datasets are available from the corresponding author upon request. Correspondence and requests for materials should be addressed to T.D.G. Matlab scripts used for the analysis of the fiber photometry data are provided at GitHub: https://github.com/Poly-03/FP-SI-code-analysis

---

**Extended data Fig 1:**
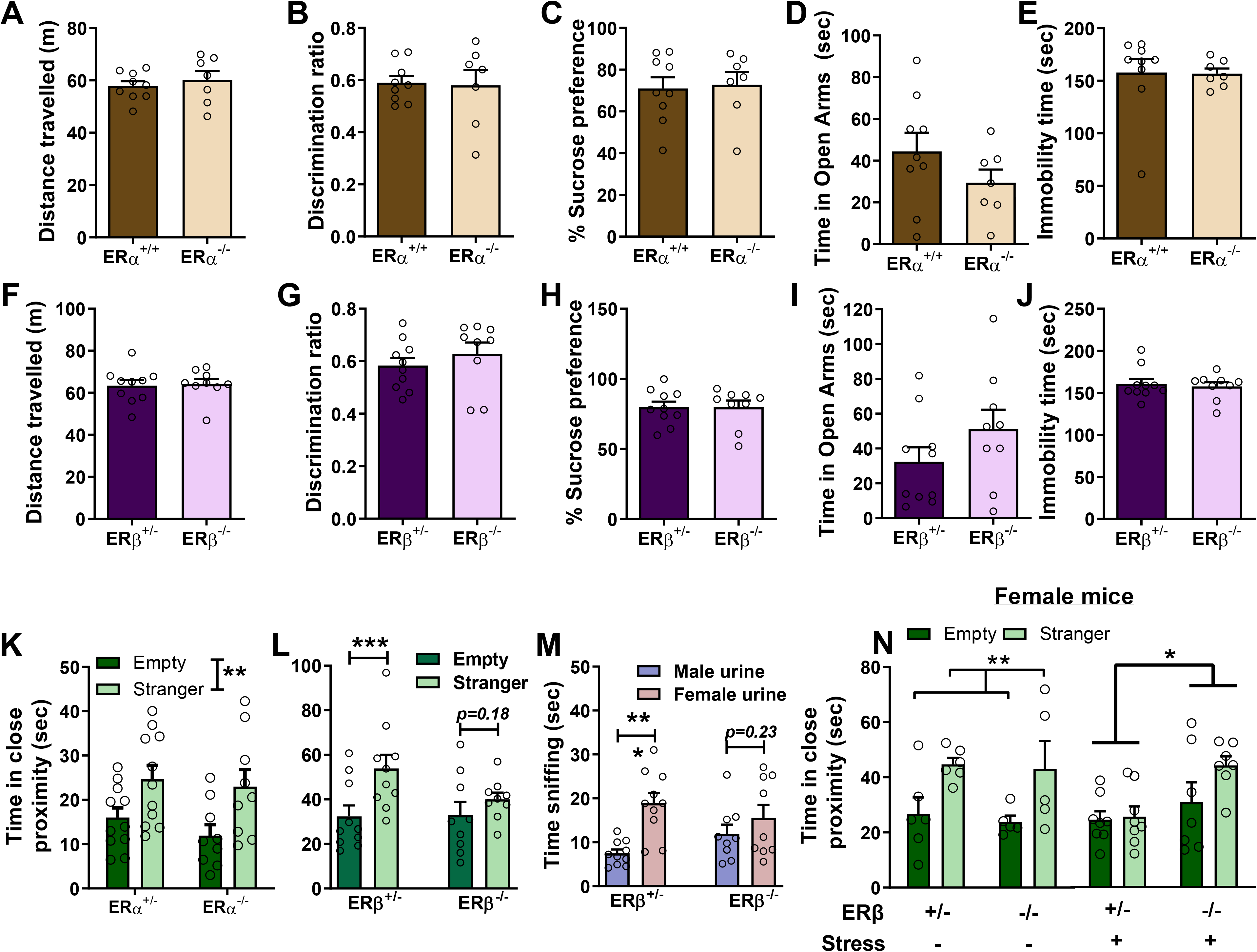
Behavioral characterization of estrogen receptor beta (ERβ) and alpha (ERα) knockout mice. Male ERα knockout mice did not demonstrate any baseline maladaptive behaviors in (A) open-field, (B) novel-object recognition, (C) sucrose preference, (D) elevated plus-maze and (E) forced-swim tests. Male ERβ knockout mice did not demonstrate any baseline maladaptive behaviors in the (F) open-field, (G) novel-object recognition, (H) sucrose preference, (I) elevated plus-maze and (J) forced-swim tests. (K) Inescapable footshock stress did not induce social interaction deficits in male ERα knockout mice. However, following footshock stress, male ERβ knockout mice developed (L) social interaction deficits and (M) anhedonia. In contrast, female ERβ knockout mice, did not reveal any social interaction deficits following inescapable footshock stress. Data shown are the mean ± S.E.M. * *p*<0.05, Abbreviations: estrogen receptor alpha, ERα; estrogen receptor beta, ERβ

**Extended data Fig 2:**
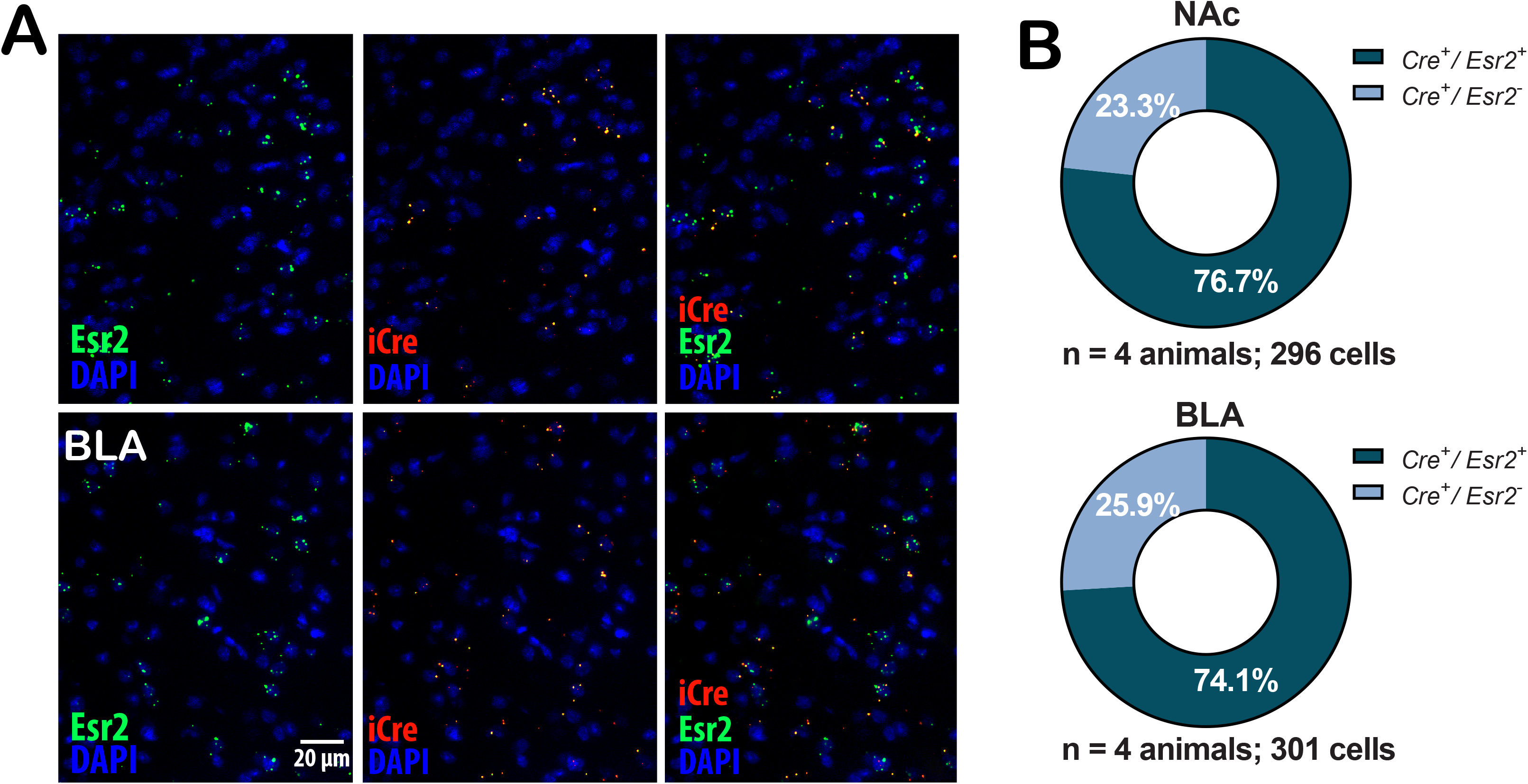
Confirmation of cre expression in ERβ expressing cells in male ERβ-cre mice. (A) Representative images and (B) quantification of RNAscope for *Esr2* and cre in DAPI-labelled cells from the NAc and the BLA in male heterozygous ERβ mice. To confirm predominant cre expression on ERβ cells, brains from male heterozygous ERβ mice were fresh frozen and processed for RNAscope. Analysis for the expression of cre on *Esr2*^+^ cells was performed. Abbreviations: basolateral amygdala, BLA; nucleus accumbens, NAc

**Extended data Fig 3:**
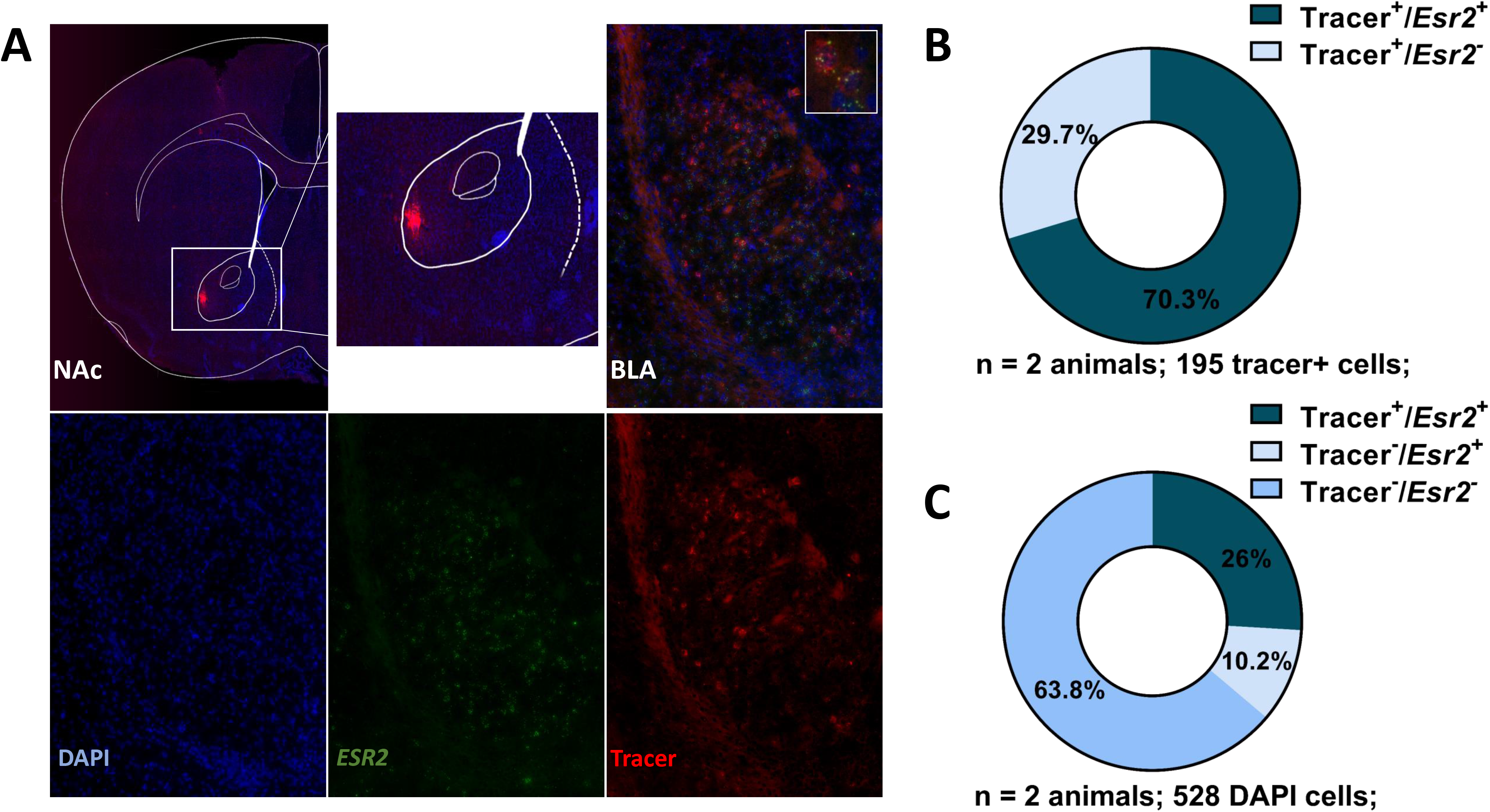
Assessment of ERβ-expressing basolateral amygdala (BLA) to nucleus accumbens (NAc) circuit. (A) Representative images, (B) quantification of RNAscope for *Esr2* (transcript coding for ERβ) and retrograde tracer positive cells and (C) quantification of RNAscope for *Esr2* (transcript coding for ERβ) and retrograde tracer positive cells and total BLA cells. Male wildtype mice received a conjugated cholera toxin (tracer) in the NAc. Following 2 weeks, the brains were fresh frozen and processed for *Esr2* expression and colocalization with tracer^+^ cells. Abbreviations: basolateral amygdala-,BLA; nucleus accumbens, NAc

**Extended data Fig 4:**
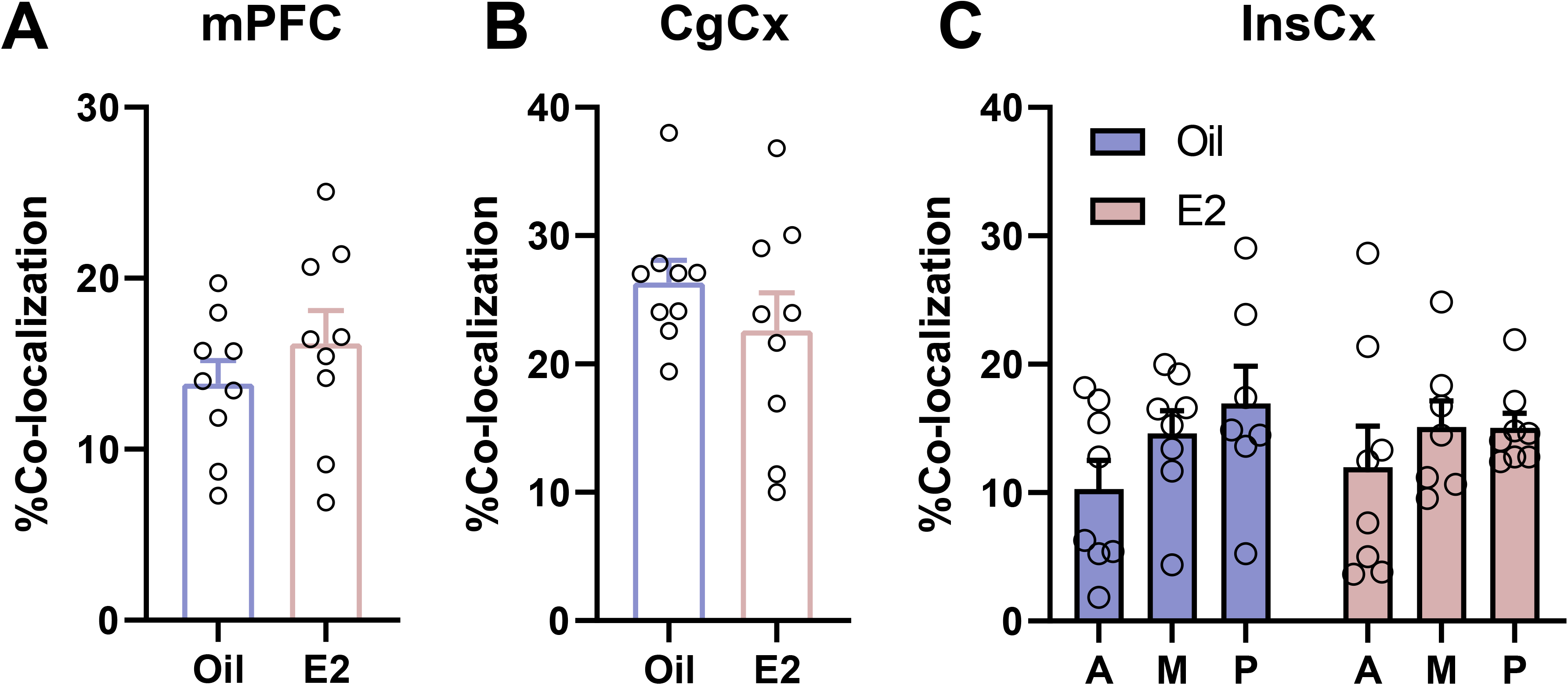
Involvement of other ERβ-projecting neurons to nucleus accumbens (NAc) in stress susceptibility. rAAV-DIO-tdTomato was injected in the NAc of male ERβ-cre mice and retrogradely transported to cell bodies localized in the BLA. Orchiectomized (ORX) mice received estradiol (E2; s.c.) 45 min prior to subthreshold social defeat stress (SSDS). No difference in the activation of the ERβ-projecting neurons to the NAc was observed from (A) medial prefrontal cortex (PFC), (B) Cingulate (CgCx) and (c) Insular cortex (InsCx).

**Extended data Fig 5:**
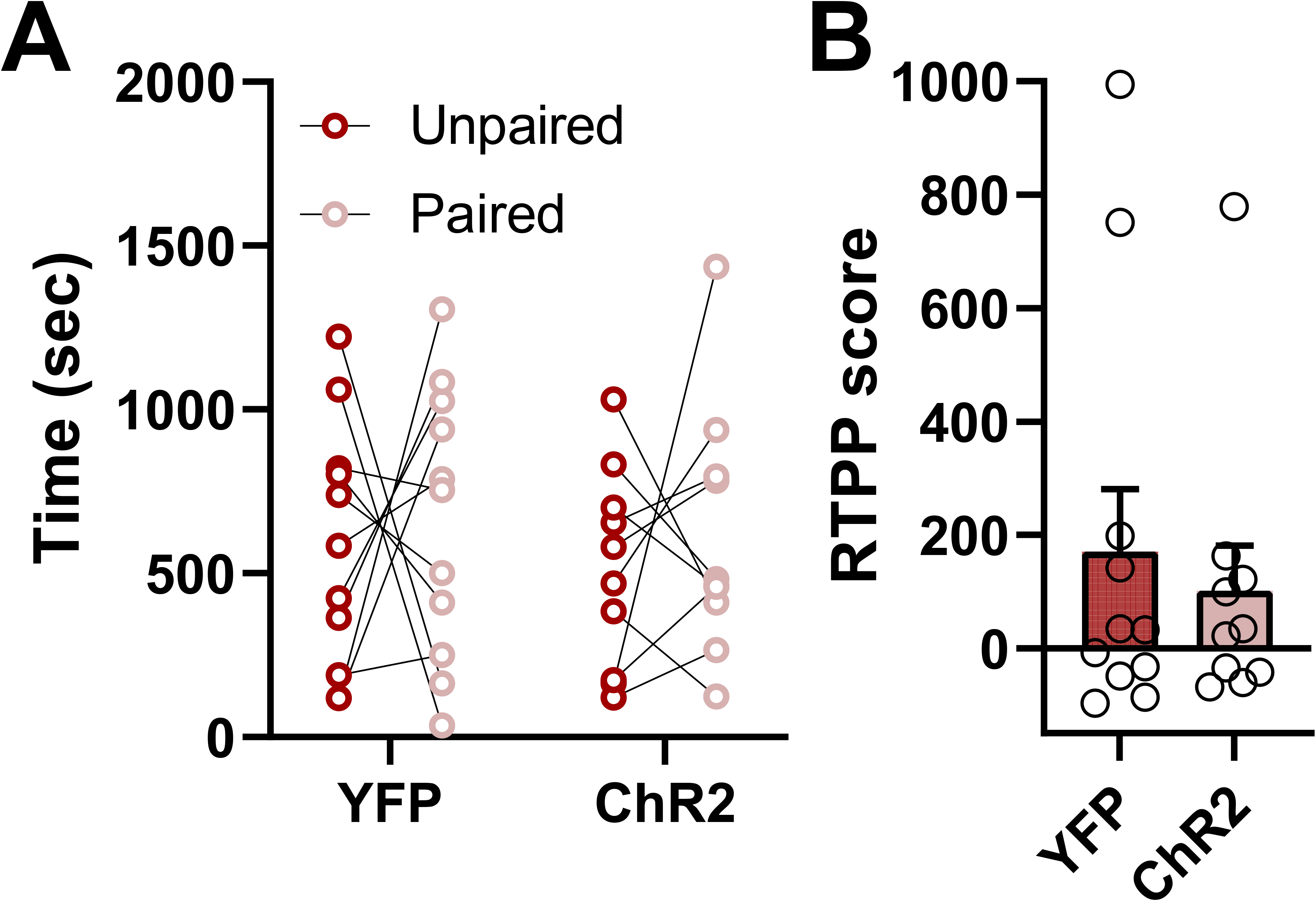
Assessment of ERβ-expressing basolateral amygdala (BLA) to nucleus accumbens (NAc) circuit in reward behaviors in female mice. ChR2-DIO-eYFP or DIO-eYFP was injected in the BLA and optic fiber was implantated in the NAc. (A-B) Blue light stimulation (470 nm; 20 Hz) of ERβ-expressing BLA to NAc terminals did not induce a real-time place preference, since we observed no difference in the time spent in the stimulation-paired compartment.

**Extended data Fig 6:**
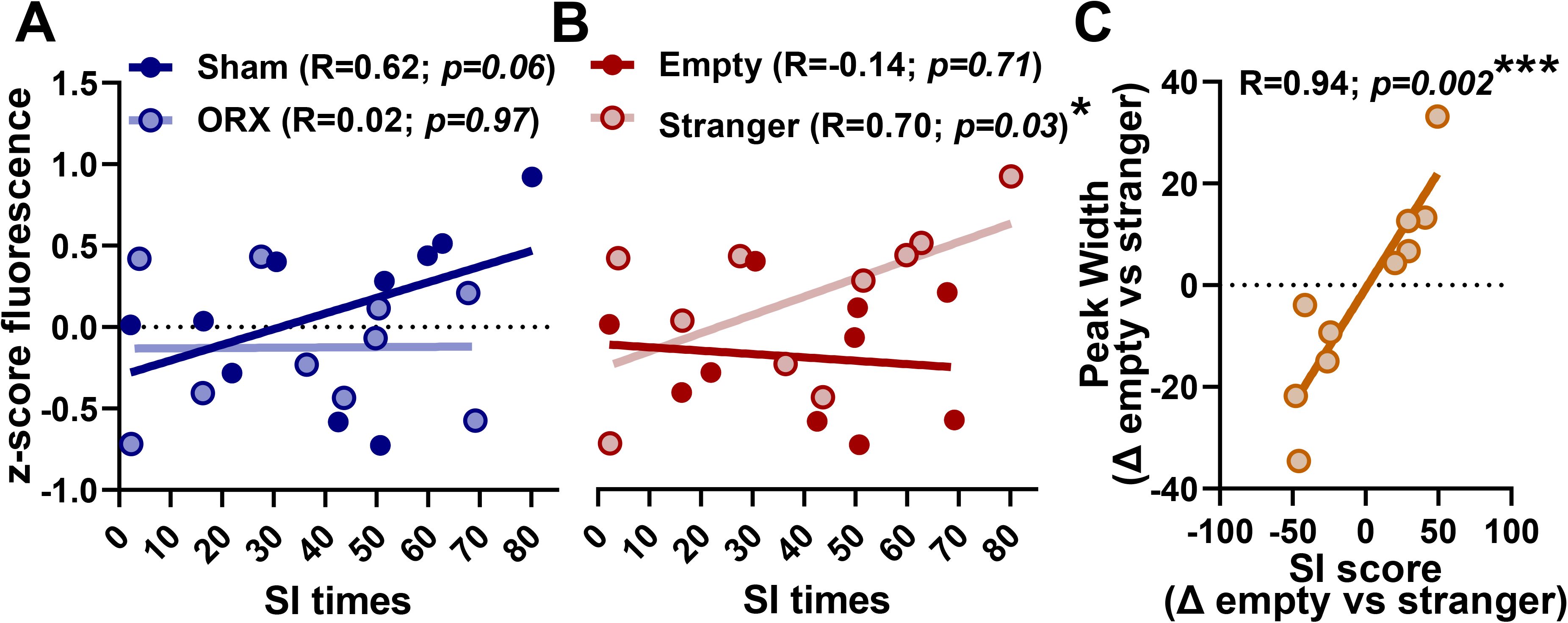
Correlations of calcium activity recorded in the nucleus accumbens terminals of the basolateral amygdala ERβ projection neurons and social interaction. Pearson’s correlation coefficients analysis demonstrated that (A) there is a statistical trend between social interaction times from gonadally intact animals and z-score fluorescence, (B) a significant correlation between interaction with the stranger mouse and z-score fluorescence and (C) a significant correlation between peak width changes and social interaction score changes. * *p*<0.05; \*\*\**p*<0.001. Abbreviations: social interaction, SI.

**Extended data Fig 7:**
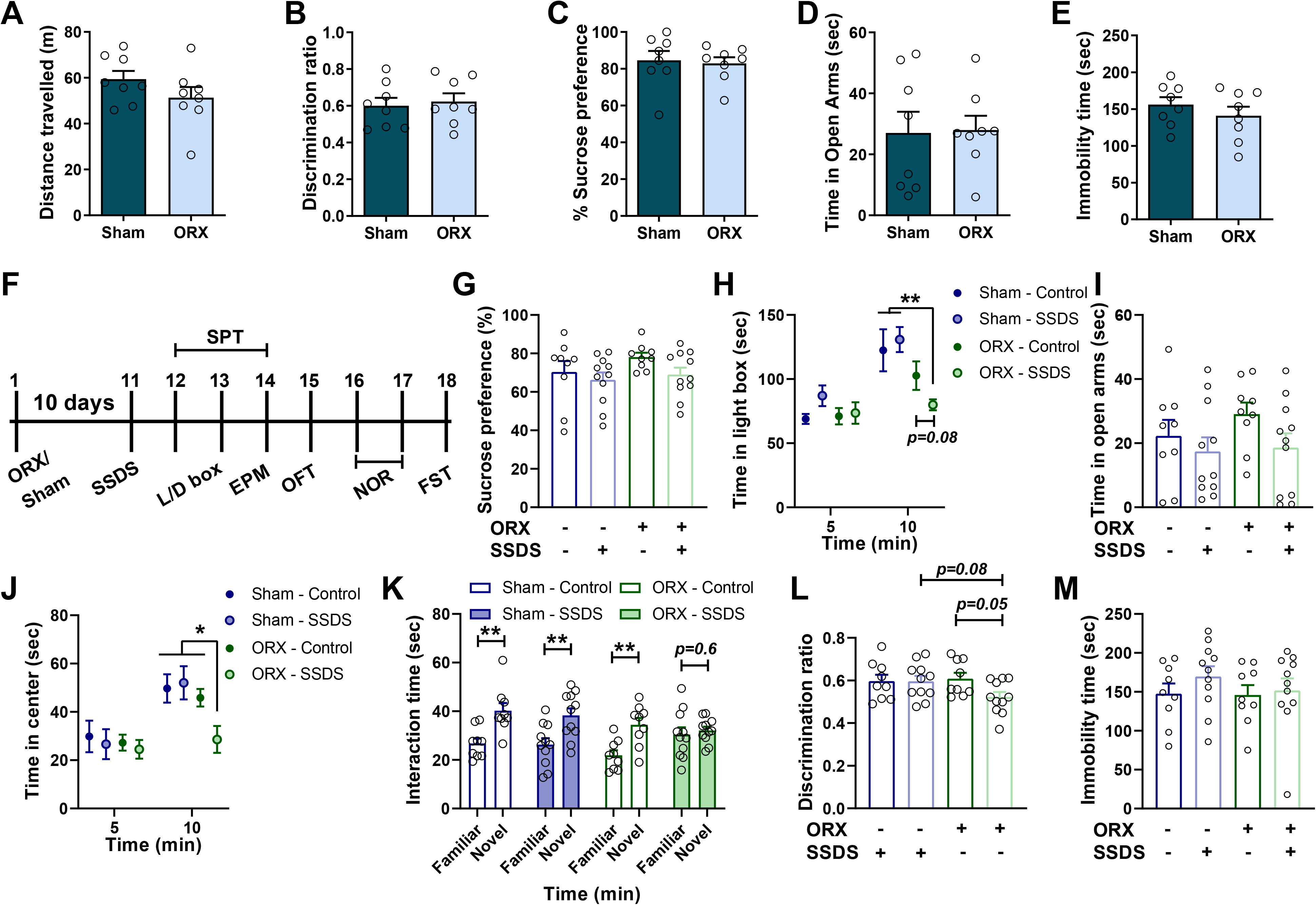
Behavioral characterization of orchiectomized (ORX) male mice. ORX mice did not demonstrate any baseline maladaptive behaviors in (A) open-field, (B) novel-object recognition, (C) sucrose preference, (D) elevated plus-maze and (E) forced-swim tests. (F) Following acute stress ORX mice underwent a battery of tests. (G) ORX mice did not show sucrose preference deficits; however, they demonstrated deficits in (H) light/dark box. (I) No difference was observed in the elevated -plus maze. (J) Decreased time spent in center in the open field test and (K-L) deficits in novel object recognition were observed. (M) No differences were found in the Forced-sim test. * *p*<0.05; \*\**p*<0.01.

**Extended data Fig 8:**
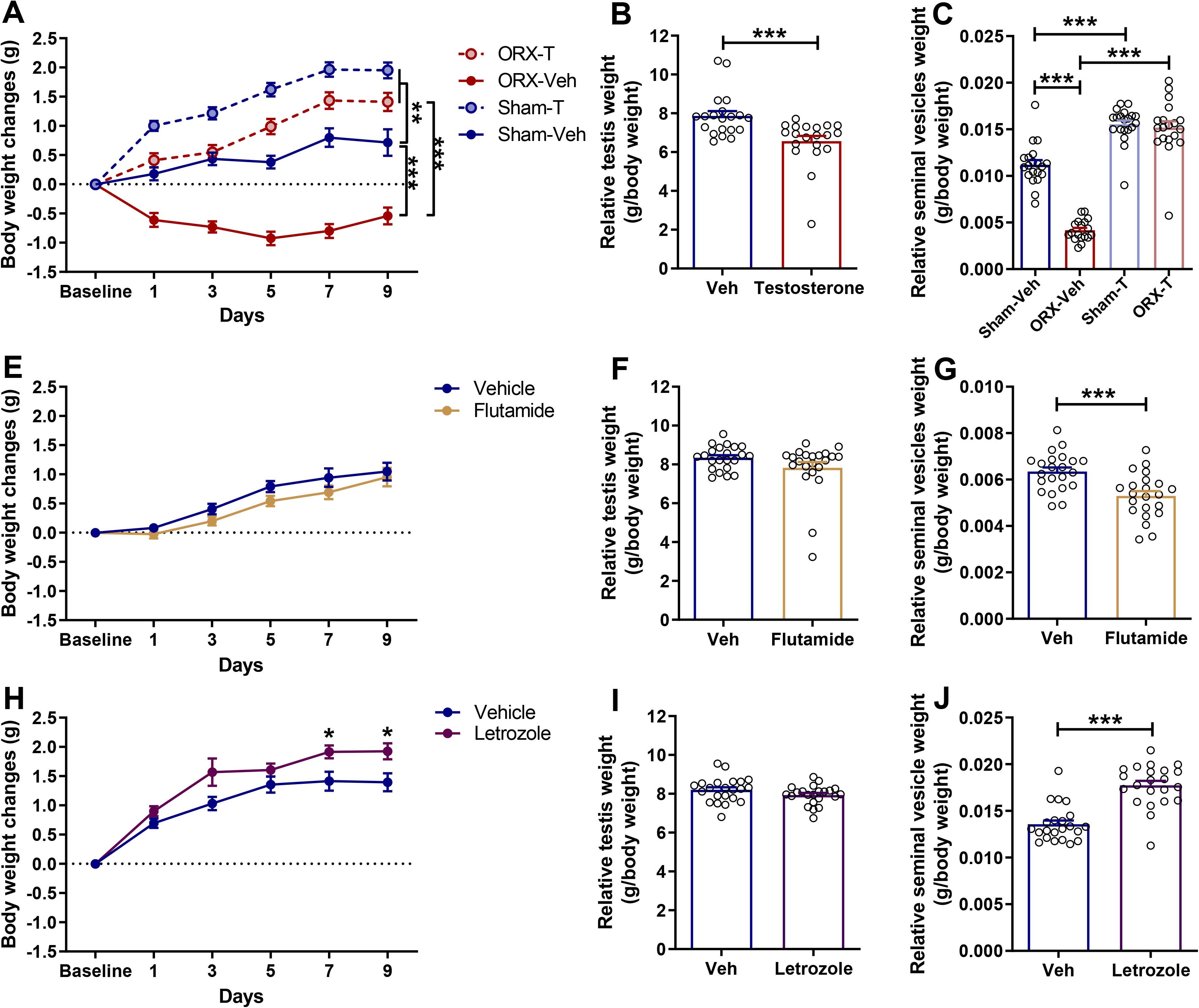
Determination of the effectiveness of testosterone, flutamide and letrozole treatments. Mice that received chronic testosterone replaced therapy showed (A) increased body weight and (B) decreased testis weight. (C) Seminal vesicles weight was decreased in orchiectomized (ORX) mice and was reversed with the chronic testosterone replacement therapy. Mice that chronically received the androgen receptor antagonist flutamide did not manifest any differences in (E) body weight or (F) testis weight. (G) Chronic treatment with flutamide resulted in a decrease in seminal vesicle weight. Mice that chronically received the aromatase inhibitor, letrozole, showed (H) increased body weight, (I) no difference in testes weight, and (J) increased seminal vesicle weight. Data shown are the mean ± S.E.M. * *p*<0.05; ** *p*<0.01; \*\*\**p*<0.001. Abbreviations: orchiectomized-,ORX; testosterone, T; vehicle, veh

**Extended data Fig 9:**
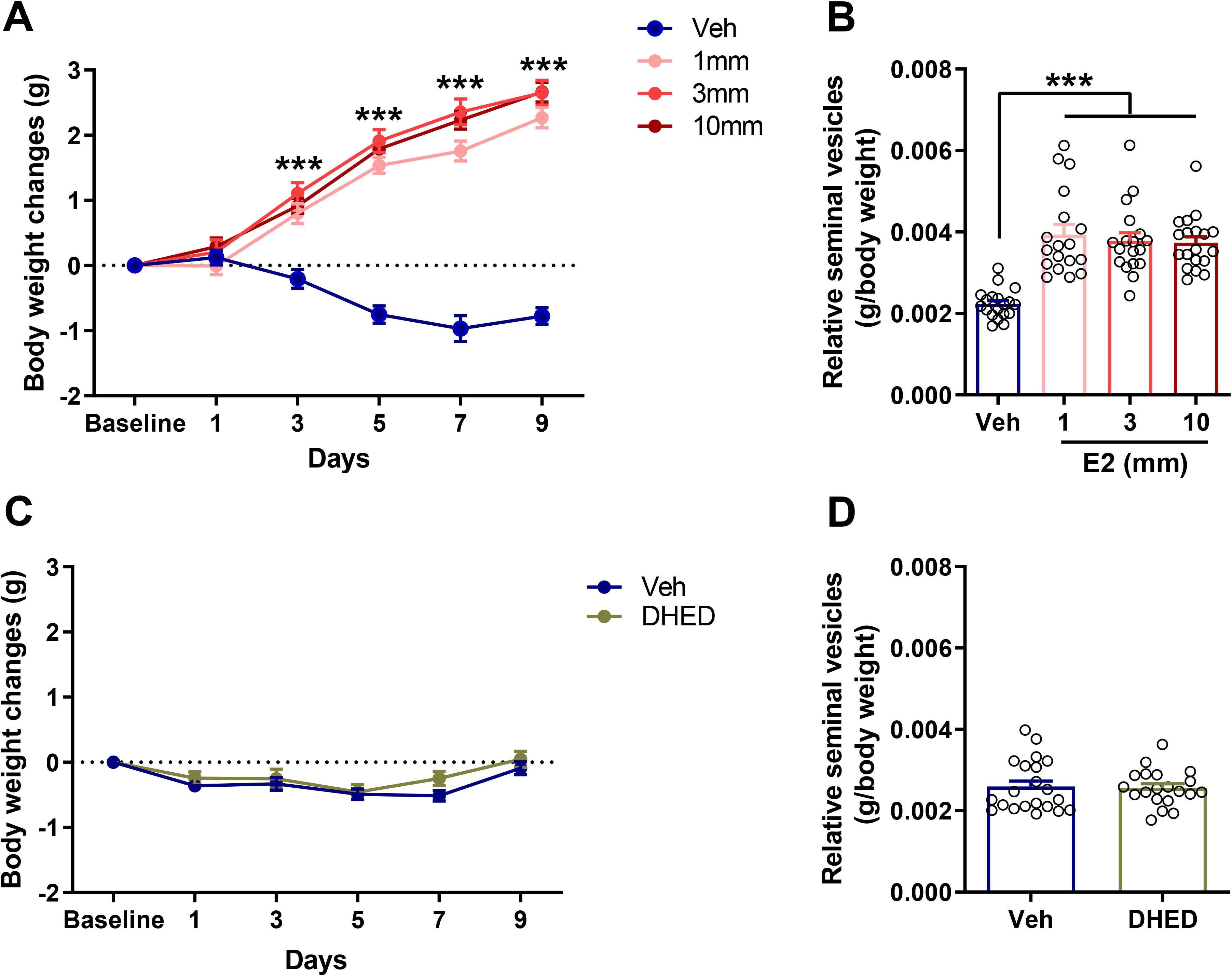
Effects of estradiol (E2) and DHED treatments on body and seminal vesicles weight changes. Mice that received estradiol (E2) treatment manifested increased (A) body weight and (B) seminal vesicles weight. Mice that received DHED treatment had no difference in their (C) body weight and (D) seminal vesicles weight compared to control mice. Abbreviations: (17β)-10,17-Dihydroxy-estra-1,4-dien-3-one, DHED; 17β-Estradiol, E2; vehicle, veh

**Extended data Fig 10:**
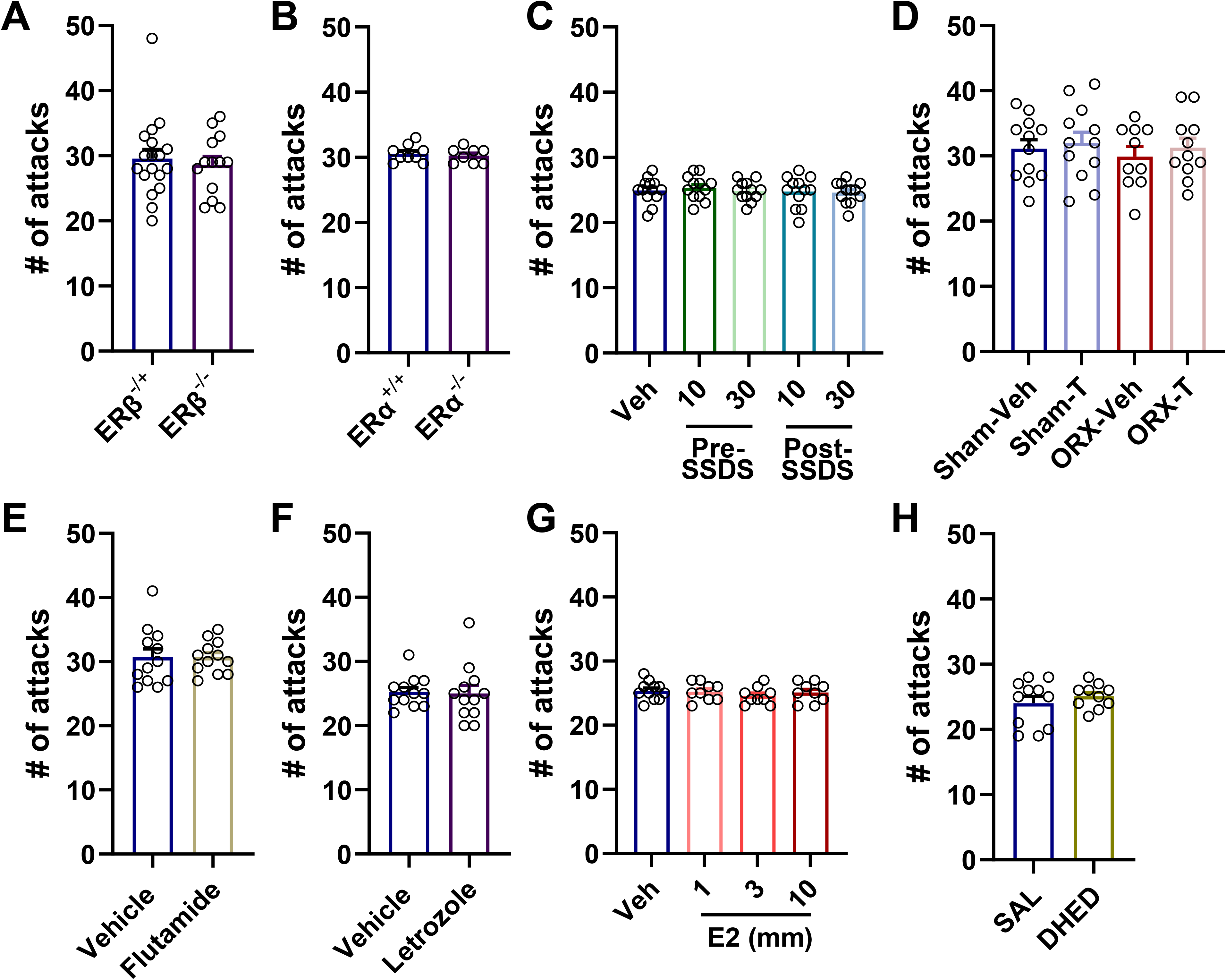
Effects of hormonal treatments on aggressiveness behaviors. No difference in the number of attacks was observed in (A) ERβ^−/−^, (B) ERα^−/−^, (C) acute E2, (D) Chronic testosterone, (E) flutamide, (F) Letrozole, (G) chronic E2, and (H) DHED treatment versus controls. Abbreviations: ( 17β)-10,17-Dihydroxy-estra-1,4-dien-3-one, DHED; 17β-Estradiol, E2; estrogen receptor Alpha, ERα; estrogen receptor beta, ERβ; testosterone, T; vehicle, veh

## Materials and Methods

### Animals

Wild-type C57BL/6J mice used for behavioral pharmacology experiments were obtained from Jackson laboratories (Bar Harbor, ME, USA). CD1 retired breeders used for the subthreshold defeat experiments and regular non-aggressive CD1 mice used for strangers in the social interaction test were obtained from Charles River (Raleigh, NC, USA). *Esr1* and *Esr2* breeding pairs were also obtained from Jackson laboratories. *Esr2*-icre knock-in (*Esr2*-icre) mice were kindly donated by Dr. CheMyong Ko (University of Illinois, Urbana-Champaign). Wild-type, heterozygous and homozygous *Esr1* knockout mice were produced in-house by breeding heterozygous males and females, while heterozygous and homozygous *Esr2* knockout mice were produced by breeding heterozygous females and homozygous males. Lastly, wild-type, heterozygous and homozygous *Esr2-*icre mice were produced in-house by breeding heterozygous or homozygous males and heterozygous females. All mouse lines were bred on a C57BL/6J background. At the time of behavioral testing, the age of the animals was in between 8 to 15 weeks. Mice were group housed and maintained under a 12 h light–dark cycle (lights on at 7:00 a.m.). Water and food were available *ad libitum*. All mice were housed in the same room in individually ventilated cages, with 3 to 5 mice per cage. All experimental procedures were approved by the University of Maryland Animal Care and Use Committee and conducted in full accordance with the National Institutes of Health Guide for the Care and Use of Laboratory Animals. Tail samples were obtained prior to weaning and genotyped by TransnetYX, Inc. (Cordova, TN, USA). All experiments and analyses were conducted in a randomized, and blind manner to avoid potential biases.

### Drugs

For the chronic administration paradigms, silastic tubes (1.02 mm inner diameter x 2.16 mm outer diameter) were filled with testosterone (10mm) or 17β-Estradiol (E2; 1,3 and 10mm; Sigma-Aldrich, St. Louis, MO, USA). Silastic implants were primed in 0.9% saline at 37^○^C overnight prior to their subcutaneous use. Letrozole (1 mg/kg/day; Sigma-Aldrich, St. Louis, MO, USA) was dissolved in propylene glycol (MP Biomedicals, Santa Ana, CA, USA) and administered through ALZET minipumps (Model 1002, 0.25μl/hour, 14 days; Cupertino, CA, USA). DHED (synthesized as previously described (*36*)) was dissolved in 0.9% saline and administered through ALZET minipumps (Model 1002, 0.25μl/hour, 14 days; Cupertino, CA, USA) at a dose of 5 μg/kg/day. Flutamide was administered through 1.5mg slow-release subcutaneous pellets (Innovative Research of America, Sarasota, FL, USA). For acute administration, 17β-Estradiol (E2; 10 and 30 μg/kg) was dissolved in sesame oil (4ml/kg, s.c.; Sigma-Aldrich, St. Louis, MO, USA).

### Surgical procedures

In all surgical procedures, mice were anesthetized with isoflurane at 3.5% and maintained at 2– 2.5% throughout the surgery. Analgesia was provided to the mice in the form of carprofen (5 mg/kg, s.c.; Norbrook laboratories, Newry, UK) prior the start of surgery and after the surgery once per day for two days.

#### Orchiectomy

Prior to incision, the scrotal area was thoroughly cleaned with Betadine solution (10% povidone-iodine surgical scrub) followed by ethanol (70%). A small cut was made in the midline of the scrotum using a pair of small sterile surgical scissors. Testicular arteries were crushed with a hemostat and cut below the crush point. Testes were separated from their blood supply, and subsequently removed. The skin was closed through suturing. Animals were allowed to recover for a minimum of ten days prior to any further experiments.

#### Subcutaneous implantation

Subcutaneous implantations were performed during the same time as the orchiectomies. The surgical area was thoroughly cleaned with Betadine solution (10% povidone-iodine surgical scrub) followed by 70% ethanol. A small, approximately 1-cm incision was made between the scapulae using a pair of small sterile surgical scissors. Hemostats were then inserted and spread laterally to the incision to create a cavity in the subcutaneous space in which ALZET mini-osmotic pumps or silastic implants were inserted for continuous drug/vehicle delivery. The skin was closed with suturing.

#### Stereotaxic surgery and viral delivery

Heterozygous *Esr2*-icre or wild-type mice were used for these experiments. Mice were positioned in a small animal stereotaxic frame (Kopf Instruments, Tujungam CA, USA) using head bars. Measurements were taken to ensure that the scalp was flat prior to recording the coordinates. All injections were performed at a rate of 20nl/min using a 1μl Hamilton Syringe (Knurled Hub Needle, 25G).

For the tracing studies, wildtype or *Esr2*-icre mice received a 350nl bilateral injection of retrograde AAV9 pCAG-FLEX-tdTomato-WPRE (Addgene,Watertown, MA, USA) or 1% retrograde conjugated cholera toxin labeled with Alexa Fluor 555 (ThermoFischer Scientific, Walthman, MA, USA) in the nucleus accumbens (AP: +1.6, ML: 1.5, DV −5.0 from the top of the skull). For the optogenetic (AAV5.EF1a-DIO-eYFP-WPRE-hGH or AAV5.EF1a-DIO-hChR2(H134R)-eYFP-WPRE-hGH; Penn Vector Core and Addgene) and fiber photometric (AAV5.Syn-Flex-GCaMP6S-WPRE-SV40; Addgene) experiments, mice received 400-nl bilateral injections in the basolateral amygdala (AP: −1.34, ML: 3.0, DV −5.5 from the top of the skull). Three weeks after viral injections, ceramic optic cannulas (400μm, 0.39 numerical aperture (NA) for optogenetics and 200mm, 0.37NA for fiber photometry; Neurophotometrics Ltd, San Diego CA, USA) were implanted in the nucleus accumbens. The use of *Esr2*-icre-mice with cre-induced expression (DIO or FLEX) virus provide specificity to ERβ expressing cells for all of our optogenetic and fiber photometry experiments. To further increase the specificity of our manipulations on ERβ-projections from the BLA to NAc, we have implanted the optic cannulas to the NAc to stimulate/record from the neuronal terminals.

Following the completion of experiments, mice were perfused with 4% paraformaldehyde; their brains were extracted and post-fixed in the same solution for at least 24 hours prior to sectioning with a vibratome (50μm thickness). Slices were then mounted, coverslipped with Vectashield Antifade Mounting Medium with DAPI (Vector Laboratories, Burlingame, CA, USA), and examined on a microscope (Leica Microsystems, DM6, Buffalo Grove, IL, USA). Fluorescent signal intensity in the GFP channel was used to confirm viral expression and the anatomical co-localization with DAPI staining was used to verify the position of optic fiber implantation.

### Behavioral experiments

#### Subthreshold social defeat stress

Retired CD1 breeders were singly housed and used as aggressors. Two days prior to the start of the experiment, experimental mice were also singly housed. The stress procedure was performed as previously described (*38*). Briefly, experimental mice were introduced to the home cage of an aggressor for 2 min to initiate the physical attack phase. Afterward, mice were transferred and housed on the opposite side of the aggressor, in the same cage separated by a perforated Plexiglass divider for 15 mins in order to maintain sensory contact. This process was repeated 3 times over a period of ~ 50 min. Following the completion of the third cycle, mice were placed back in their home cages.

Acute E2 was administered 45 min prior to the initiation of stress or 5 min after the completion of stress. For the optogenetic stimulation, ChR2 and YFP-injected mice received a 4 Hz, 5 ms pulse width, 10-15 mW 470 nm light stimulation for 45 min prior to the initiation of stress. This stimulation paradigm was shown previously to increase synaptic transmission (*19*). The optogenetic set up consisted of an LED driver paired with a fiber-coupled LED light source attached to a bifurcated optogenetic path cable (400μm, 0.49NA; ThorLabs, Newton NJ, USA).

#### Inescapable foot-shock stress

Experimental mice were placed in one side of two-chambered shuttle boxes (Coulbourn Instruments, Whitehall, PA, USA), with the door between the chambers closed. Following a 5-min adaptation period, mice received foot shocks (0.3 mA, 2-sec shock duration, 15-sec inter-trial interval) for 51 min which is equivalent to the duration of the subthreshold social defeat stress.

#### Social Interaction

Social interaction was performed 24 hours following subthreshold social defeat or inescapable foot-shock stress. For the habituation phase, mice were placed in a rectangular box (40 cm length × 30 cm width × 35 cm height; Stoelting, IL) divided into three compartments (two equal-sized end-chambers and a middle chamber) for five minutes (10-15 lux). After the habituation phase, two small wire cages were introduced into the two end chambers, one containing an unfamiliar, non-aggressive CD1 mouse and the other remaining empty. The amount of time mice spent sniffing each cage during the five-min test was assessed using CleverSys tracking software (CleverSys, Inc, Reston VA, USA).

#### Female Urine Sniffing Test (FUST)

FUST procedure was performed 24 hours after the social interaction test and 48 hours after stress. FUST was performed as previously described (*39*). The amount of time mice spent interacting with a cotton-tipped applicator soaked in either fresh male or female mouse urine during a total of 3 minutes was analyzed by an experimenter blind to the experimental groups.

#### Open-Field Test (OFT)

OFT was performed under 300-Lux white light. Mice were individually placed into open-field arenas (100 × 100 × 38 cm; San Diego Instruments, San Diego, CA) for a 10-min period. The sessions were recorded using an overhead, digital video camera. Distance traveled and time spent in the center of the arena was analyzed using TopScan v2.0 (CleverSys, Inc., Reston VA).

#### Novel-Object Recognition (NOR)

Short-term recognition memory was assessed using the novel object recognition task protocol, as previously described (*16*). NOR was performed in dim white lighting conditions (~10–15 lux). The apparatus and objects used here has been previously described (*40*). The test was conducted over two days. On the first day, the habituation phase, the animals explored an empty NOR apparatus (40 × 9 × 23 cm) for 30 min and then returned to their home cages. On the second day, the mice were re-introduced into the same apparatus, but this time containing two identical objects fixed onto the floor, which they explored for 30 min. After this familiarization phase, mice were immediately returned to their home cages for another 30 min. The mice were then placed back into the NOR apparatus, in which one of the “familiar” objects was replaced by a “novel” object for a 4 min test phase. All three phases of NOR were recorded via an overhead video camera and analyzed using TopScan v2.0 automated scoring software (CleverSys, Inc., Reston VA). The time spent interacting with the familiar and novel objects during the retention phase was measured. A discrimination ratio was calculated by dividing the time spent interacting with the novel object by the total time spent interacting with both objects during the retention phase.

#### Light-Dark box

The Light-Dark box was used as previously described (16). Briefly, mice were placed in the illuminated compartment of the L/D box (35 × 35 cm), facing the wall opposite to the dark compartment, and allowed to explore the whole apparatus for 5 min. The sessions were recorded using a video-camera and the time spent in the illuminated and dark compartment was scored using TopScan v2.0 (CleverSys, Inc., Reston VA).

#### Sucrose preference test

Mice were singly housed and presented with two identical bottles containing either tap water or 1% sucrose solution. The sucrose consumption was measured over 48 hours. The location of the sucrose and tap water bottles was changed every day to avoid the development of side preference.

#### Elevated-Plus Maze (EPM)

EPM was carried out in dim white lighting conditions (~5 lux). The EPM apparatus consisted of 2 closed arms and 2 open arms (39 × 5 cm each) and was elevated 50 cm above the floor (Stoelting, Woodale, IL). The experiment was carried out as previously described ^5^. The time spent in the open and closed arms of EPM during the 5-min test was recorded by an overhead digital video camera and scored using TopScan v2.0 (CleverSys, Inc., Reston VA). Amount of time spent in the open arms was used as the primary outcome for the anxiety behavioral assessment.

#### Forced-Swim Test (FST)

FST was performed in normal white light conditions (~300 lux) as previously described (*41*). Briefly, mice were subjected to a 6-min swim session in clear Plexiglass cylinders (30-cm height × 20-cm diameter) filled with 15 cm of water (23 ± 1°C). Sessions were recorded using a digital video camera. Immobility time, defined as passive floating with no additional activity other than that necessary to keep the animal’s head above water, was scored for the last 4 min of the 6-min test by a trained experimenter blind to the genotypes.

#### Real-time place preference

The same three-chambered box or the social interaction test was used. However, for this test, one compartment was altered with white vertical stripes 4 cm apart on each wall and a smooth grey floor, while the other with white horizontal stripes 4 cm apart on the walls and a smooth grey floor. Allocation of the light-paired compartment was counterbalanced to avoid any side preferences. ChR2 and YFP-injected mice were connected to optogenetic fiber cables (ThorLabs, Newton NJ, USA) and placed in the middle compartment, and tracked by an overhead camera. During the 30 min testing period, mice who crossed into the allocated light-paired compartment received a 20Hz continuous 470nm ~10-15mW stimulation. Mouse crossings into each compartment were detected by a Bonsai script which communicated to the LED through connected Arduinos for light delivery.

### Whole-cell voltage-clamp electrophysiology

Mice (n=3 males) that were previously injected with ChR2 in the BLA were deeply anesthetized with isoflurane before decapitation and brain extraction. Coronal 250 μm thick sections were collected in ice cold 95% oxygen, 5% carbon dioxide-bubbled modified artificial cerebrospinal fluid (aCSF, 194mM sucrose, 30mM NaCl, 4.5mM KCl, 1mM MgCl2, 26mM NaHCO3, 1.2mM NaH2PO4, and 10mM D-glucose) before incubating at 32°C for 30 min in aCSF (124mM NaCl, 4.5mM KCl, 2mM CaCl2, 1mM MgCl2, 26mM NaHCO3, 1.2mM NaH2PO4, and 10mM D-glucose). Slices were then stored at room temperature until recording.

Slices were hemisected, placed into a recording chamber, and perfused with temperature controlled aCSF (29-31°C). NAc medium spiny neurons (MSNs) were visualized using infrared differential interference contrast light microscopy, Q-capture camera, and associated Pro 7 software. MSNs were voltage clamped at −60 mV using a MultiClamp 700B Amplifier (Molecular Devices, San Jose, CA, USA). Optically evoked post synaptic currents (oPSC) were elicited by an optical fiber placed in the recording chamber that delivered 4 ms pulses of light (473 nm) every 20s for a total recording duration of 15 min. oPSCs were recorded using borosilicate glass pipettes (2-4 MΩ resistance) filled with Cesium Methanesulfonate internal solution (135mM cesium methanesulfonate, 3mM NaCl, 10mM HEPES,0.6mM EGTA, 4mM MgATP,0.3mM NaGTP,5mM QX-314Cl, pH 7.2, 310mOsm). Signals were filtered at 2 kHz, digitized at 10 kHz, and acquired using the Clampex 10.4.1.4 software (Molecular Devices). oPSC amplitudes were averaged per minute and expressed as a percentage change from baseline measurements (5 min baseline recording). Following the baseline recording, the AMPAR blocker NBQX (5μM) and NMDAR blocker APV (50μM) were drip perfused into the recording chamber for a 10-min recording period.

### *In vivo* fiber photometry

Heterozygous *Esr2*-icre mice received bilateral infusions of AAV-FLEX-GCaMP6s in the BLA. Three to four weeks after the virus infusion, mice underwent orchiectomy or sham surgeries and were bilaterally implanted with fiber optic cannulae (200μm, 0,37NA) in the NAc. Following a minimum of 10-day recovery, mice went through subthreshold social defeat stress. Twenty-four hours after the exposure to stress, mice went through the social interaction test. On the day of the social interaction test, mice were connected to the fiber photometry system (Neurophotometrics, San Diego, CA, USA) to record calcium transients in the BLA to NAc afferent terminals using a low-autofluorescence bifurcated optic patch cable (Doric Lenses Inc., Quebec, Canada). Independence of signals was tested prior to the start of the experiment. Excitation wavelengths used were 470 nm for the calcium-dependent GCaMP6s signal, temporally interleaved with 410 nm light at 40 Hz for the calcium-independent (isosbestic) signal. Mice were placed and habituated to the three-chamber arena for measuring social interaction for 5 min. During this period, the photometer’s excitation LEDs were on to minimize any rapid initial photobleaching during the test. For the test, a stranger (novel CD1 mouse) was placed in one of the two chambers before reintroducing the test mouse. During a test period of 5 minutes, photometry data was recorded, and the initiation and termination of interactions with either the occupied (stranger) or empty chambers were manually timestamped by the experimenter.

The fiber photometry data was analyzed offline using custom-created MATLAB scripts. We also made use of code from Ekaterina Martianova (*42*). Briefly, photometry signals were deinterleaved, and then corrected for baseline drift due to photobleaching using the AirPLS algorithm by Zhang et al (*43*). A smoothing filter was applied to minimize the high-frequency noise. After z-scoring both signals, the 410 nm signal was scaled and fitted to the 470 nm one before subtraction to correct for any motion artifacts. The resulting z-scored, motion-corrected 470 nm fluorescent signal was used for all subsequent analyses. Using the manually encoded timestamps, we defined periods of interaction with either the stranger or the empty chambers and used them to further analyze the signal. For each animal, we generated a mean z-scored fluorescence response to each type of interaction by averaging the segments of the signal ranging temporally from 120 frames (5.76s) prior to the initiation of interaction to 120 frames (5.76s) after the interaction. Any interactions of a type that occurred within 5 s of each other were treated as a single interaction for this analysis, and interactions that occurred within 120 frames of the start or the end of the recording were excluded. For the time-locked analysis, a baseline correction was applied to each response prior to averaging, to account for variation in the fluorescence preceding the interaction. This involved zeroing the mean z-scored fluorescence value corresponding to the period of 90 to 40 frames before the interaction. This baseline period of 2.4 s was chosen as it did not appear to overlap with any observed “anticipatory” activity related to the interactions. Individually, the resultant mean z-scored fluorescence responses were used for the final analysis, which involved comparing the group mean AUC before and after social interaction for each condition.

### Pharmacokinetic measurements of distribution of DHED-derived estradiol

The experimental design (including the synthesis of a deuterium-labelled DHED) was adapted from a previously described study ^1^. Briefly, mice (n=3-4/timepoint) received d3-DHED (200 μg/kg, s.c.,) and euthanized 0.5, 1, 2, 4 and 8 hours after the injection. Blood was collected by cardiac puncture and the animals were then perfused with cold saline through the left ventricle, and brain tissues were harvested immediately after finishing the perfusion. The brain (cortical) tissues were homogenized in pH 7.4 phosphate buffer to obtain 20% w/v homogenates. Serum samples were obtained from the collected blood by centrifugation (1,500g at 4 °C for 10 min). Sera and brain homogenates were processed by liquid-liquid extraction using ^13^C-labeled 17β-estradiol (^13^C_6_-E2 added at 100 pg/ml serum and 1.7 ng/g wet tissue) as internal standard followed by quantitative isotope-dilution liquid chromatography–tandem mass spectrometry (LC–MS/MS) analyses after dansylation as described previously ^1^.

### cFos immunofluorescence

*Esr2*-icre mice received an injection of rAAV9 pCAG-FLEX-tdTomato-WPRE virus. Following 4 weeks, mice were either treated with E2 or Veh and underwent subthreshold social defeat stress. Two hours following the treatment injections mice were perfused with 4% paraformaldehyde. Brains were extracted, post-fixed via immersion in 4% paraformadehyde for 24 hours at 4°C, and cryoprotected in 30% sucrose solution. Consecutive 30μm-thick free-floating sections were collected in a cryostat (Leica CM3050S) and stored in cryoprotectant at −20°C until further processing for immunofluorescence.

Six representative brain sections (each 180 μm apart) containing the BLA, mPFC, CgCx and InsCx were selected for each animal. Sections were washed in 1X tris buffered saline (TBS) three times for 15 minutes each, blocked in 5% normal horse serum in 0.4% in Triton X-100 in 1X TBS for an hour at room temperature (RT), and incubated in monoclonal primary antibody against cFos (Cell Signaling Technologies 2250S, 1:3000) for an hour at RT, then 48 hours at 4°C. Sections were protected from light at all washes and incubations to prevent potential photobleaching.

Sections were then washed in 1X TBS 3 times for 15 minutes each, and incubated in Alexa fluor 488 conjugated secondary antibody (Cell Signaling Technologies 4412S, 1:2500). After secondary antibody incubation, the sections were washed in 1X TBS two times for 15 minutes each, and in 1X PBS for an additional 15 min. Finally, sections were nuclear stained in DAPI, mounted, coverslipped using Vectashield Vibrance antifade mounting medium (Vector Laboratories), and allowed to set overnight prior to imaging.

Multichannel tiled z-stacked fluorescent images were acquired using a Nikon W1 spinning disk confocal microscope with the 40X objective and following filter cubes: DAPI (405 nm), GFP (488 nm), and RFP (561 nm). All images corresponding to a single immunofluorescence experiment were taken with constant excitation laser intensity, exposure, and display adjustment. Image stitching and projection were completed using NIS-elements software (Nikon, Melville, NY, USA).

For the quantitative assessment of cFos and rAAV colocalization, images were imported into Bitplane software Imaris9.6 (Oxford instruments, Concord, MA, USA). A set of 3D reconstructed surface objects were created for the GFP and RFP channel by mapping and thresholding the fluorescent signals from cFos immunohistochemistry and endogenous tdTomato tag expressed by the virus. Next, objects were segmented into individual cells using the built-in watershed algorithm. Colocalization was then determined by setting the shortest distance between the two distance-transformed classes of objects to zero. All images corresponding to a single experiment were batch-analyzed together, with the program automatically generated the count for total number of rAAV labeled cells and the number cFos-rAAV colocalized cells. After which, a scorer blind to group assignments screened through the automatic detection results and manually confirmed its accuracy.

### RNAscope

RNAscope Fluorescent Multiplex Detection Reagent Kit v2 (ACDBio, Newark, CA, USA) was used to assess the colocalization of *Esr2* and iCre in the heterozygous *Esr2*-iCre line and the colocalization of *Esr2* to the CTb retrograde tracer targeting the BLA to NAc projections. Four heterozygous *Esr2*-iCre mice and two wildtype mice with CTb retrograde tracer injected in the NAc were briefly (~1min) anesthetized with 3.5% isoflurane. Animals were then decapitated, their brains extracted and rapidly frozen in isopentane. 20 μm thick fresh frozen brain sections were collected with a cryostat, directly mounted, and fixed in 4% paraformaldehyde in 1x phosphate buffered saline (15min), and dehydrated in graded ethanol solutions (50% 70% 100% 100%, 5min each). Sections were air dried and hydrophobic barrier pen outlines were drawn prior to treatment with hydrogen peroxide (10min) and Protease plus (5mins), with two 1xPBS washes (2min each) in between. All of the remaining incubation steps were completed in the RNAscope hybridization oven at 40ᵒC. Sections were hybridized with probes for Esr2 and iCre (50:1 dilution), or Esr2 alone for the retrograde tracer animals (2hrs). After 2 washes in RNAscope Washing Buffer (2mins each), sections were incubated with AMP1 (30min), AMP2 (30min), AMP3 (15min) with two Washing Buffer washes (2min each) in between each step. Esr2 signals were then developed with successive incubation in HRP-C1 (15min), TSA plus fluorescein (30 min, 1:1000 in TSA diluent), and HRP blocker (20min) with 2 intervening washes in RNAscope Washing Buffer (2min each). For the heterozygous ERᵦ-iCre mice, iCre signals were developed by successive incubation in HRP-C2 (15 min), TSA plus Cy3 (30min, 1:1000 in TSA diluent), and HRP blocker (20min) with 2 intervening washes with RNAscope Washing Buffer between each step. Upon completion of signal development, all sections were stained with DAPI for 5min, coverslipped with FluorSave mounting medium, and set overnight at 4ᵒC prior to imaging.

BLA and NAc sections were imaged at 20x for all animals. Nuclei expressing ≥2 foci of both Esr2 and iCre were counted as colocalization. These colocalized cells were contrasted to the total number of cells expressing ≥2 foci of iCre to determine the percentage of colocalization of Esr2 to iCre.

The colocalization of Esr2 to a retrograde tracer targeting the projections from the BLA to NAc was similarly determined through counting the total number of nuclei expressing retrograde tracer and ≥2 foci of Esr2. The total colocalized *Esr2* + tracer nuclei were contrasted to the total number of nuclei expressing tracer (both alone and when colocalized) to determine the percentage of colocalization of Esr2 to tracer. All the images obtained from the RNAscope were manually analyzed by an experienced experimenter blind to the experimental groups.

### Statistical Analysis

Details of all n numbers. statistical analyses and results are provided in the Supplementary Table 1. Required samples sizes were estimated based upon on our past experience performing similar experiments. All studies were performed using simple and stratified randomization methods. Studies were performed using distinct animal groups with the exception of the baseline behaviors where a battery of behavioral experiments was performed using the same mice. Experimentation and analyses were performed by experimenters’ blind to the group assignments. All statistical tests were two-tailed, and significance was set at p<0.05. Data were tested for equal variance and Gaussian distribution using the Brown-Forsythe test and Shapiro-Wilk test, respectively using SigmaPlot v14.5. When normality and equal variance was achieved, parametric statistical tests were performed. When normality of samples failed non-parametric tests were used and when equal variance failed Brown Forsythe ANOVA test was used. In the cases were there were more than 2 levels of repeated or matched variables with ∊<1.00, Geisser Greenhouse correction was used. Planned comparisons were performed in the cases that there was a priori hypothesis. Correction for multiple comparisons following ANOVAs, Kruskal-Wallis, Brown Forsythe ANOVA test, was performed using either Holm-Sidak post-hoc test or the two-stage linear step up procedure of Benjamini, Krieger and Yekutieli. Spearman correlation analysis was used to test relationships between fiber photometry signals and behavioral outcomes. Statistical analyses were performed using GraphPad Prism v9.0 and SigmaPlot v14.5.

## References and Notes

1. G. V. Callard, Z. Petro, K. J. Ryan, Conversion of Androgen to Estrogen and Other Steroids in the Vertebrate Brain. American Zoologist 18, 511–523 (2015).

2. G. Perez-Palacios, K. Larsson, C. Beyer, Biological significance of the metabolism of androgens in the central nervous system. J Steroid Biochem 6, 999–1006 (1975).

3. P. B. Danielson, The cytochrome P450 superfamily: biochemistry, evolution and drug metabolism in humans. Curr Drug Metab 3, 561–597 (2002).

4. W. Wharton, C. E. Gleason, S. R. Olson, C. M. Carlsson, S. Asthana, Neurobiological Underpinnings of the Estrogen - Mood Relationship. Curr Psychiatry Rev 8, 247–256 (2012).

5. J. A. McHenry et al., Hormonal gain control of a medial preoptic area social reward circuit. Nat Neurosci 20, 449–458 (2017).

6. K. E. Yoest, J. A. Cummings, J. B. Becker, Estradiol, dopamine and motivation. Cent Nerv Syst Agents Med Chem 14, 83–89 (2014).

7. S. S. Kokane, L. I. Perrotti, Sex Differences and the Role of Estradiol in Mesolimbic Reward Circuits and Vulnerability to Cocaine and Opiate Addiction. Front Behav Neurosci 14, 74 (2020).

8. C. de Novaes Soares, O. P. Almeida, H. Joffe, L. S. Cohen, Efficacy of Estradiol for the Treatment of Depressive Disorders in Perimenopausal Women: A Double-blind, Randomized, Placebo-Controlled Trial. Archives of General Psychiatry 58, 529–534 (2001).

9. M. A. Whooley, D. Grady, J. A. Cauley, Postmenopausal estrogen therapy and depressive symptoms in older women. J Gen Intern Med 15, 535–541 (2000).

10. P. J. Schmidt et al., Estrogen replacement in perimenopause-related depression: a preliminary report. Am J Obstet Gynecol 183, 414–420 (2000).

11. S. N. Seidman, J. G. Rabkin, Testosterone replacement therapy for hypogonadal men with SSRI-refractory depression. J Affect Disord 48, 157–161 (1998).

12. M. M. Shores, D. R. Kivlahan, T. I. Sadak, E. J. Li, A. M. Matsumoto, A randomized, double-blind, placebo-controlled study of testosterone treatment in hypogonadal older men with subthreshold depression (dysthymia or minor depression). J Clin Psychiatry 70, 1009–1016 (2009).

13. A. Walther, J. M. Wasielewska, O. Leiter, The antidepressant effect of testosterone: An effect of neuroplasticity? Neurology, Psychiatry and Brain Research 32, 104–110 (2019).

14. S. lorA. Golden, H. E. Covington, 3rd, O. Berton, S. J. Russo, A standardized protocol for repeated social defeat stress in mice. Nat Protoc 6, 1183–1191 (2011).

15. O. Malkesman et al., The female urine sniffing test: a novel approach for assessing reward-seeking behavior in rodents. Biol Psychiatry 67, 864–871 (2010).

16. P. Georgiou, P. Zanos, C. E. Jenne, T. D. Gould, Sex-Specific Involvement of Estrogen Receptors in Behavioral Responses to Stress and Psychomotor Activation. Front Psychiatry 10, 81 (2019).

17. A. E. Kudwa, C. Bodo, J. A. Gustafsson, E. F. Rissman, A previously uncharacterized role for estrogen receptor beta: defeminization of male brain and behavior. Proc Natl Acad Sci U S A 102, 4608–4612 (2005).

18. E. J. Nestler, Role of the Brain’s Reward Circuitry in Depression: Transcriptional Mechanisms. Int Rev Neurobiol 124, 151–170 (2015).

19. R. C. Bagot et al., Ventral hippocampal afferents to the nucleus accumbens regulate susceptibility to depression. Nat Commun 6, 7062 (2015).

20. M. Heshmati et al., Depression and Social Defeat Stress Are Associated with Inhibitory Synaptic Changes in the Nucleus Accumbens. J Neurosci 40, 6228–6233 (2020).

21. S. A. Golden et al., Epigenetic regulation of RAC1 induces synaptic remodeling in stress disorders and depression. Nat Med 19, 337–344 (2013).

22. T. C. Francis et al., Nucleus accumbens medium spiny neuron subtypes mediate depression-related outcomes to social defeat stress. Biol Psychiatry 77, 212–222 (2015).

23. J. S. Brog, A. Salyapongse, A. Y. Deutch, D. S. Zahm, The patterns of afferent innervation of the core and shell in the “accumbens” part of the rat ventral striatum: immunohistochemical detection of retrogradely transported fluoro-gold. J Comp Neurol 338, 255–278 (1993).

24. M. J. Christie, R. J. Summers, J. A. Stephenson, C. J. Cook, P. M. Beart, Excitatory amino acid projections to the nucleus accumbens septi in the rat: a retrograde transport study utilizing D[3H]aspartate and [3H]GABA. Neuroscience 22, 425–439 (1987).

25. P. O’Donnell, A. Grace, Synaptic interactions among excitatory afferents to nucleus accumbens neurons: hippocampal gating of prefrontal cortical input. The Journal of Neuroscience 15, 3622–3639 (1995).

26. M. Blurton-Jones, M. H. Tuszynski, Estrogen receptor-beta colocalizes extensively with parvalbumin-labeled inhibitory neurons in the cortex, amygdala, basal forebrain, and hippocampal formation of intact and ovariectomized adult rats. J Comp Neurol 452, 276–287 (2002).

27. T. P. O’Leary et al., Extensive and spatially variable within-cell-type heterogeneity across the basolateral amygdala. Elife 9, (2020).

28. S. G. Chrysant, G. S. Chrysant, Cardiovascular benefits and risks of testosterone replacement therapy in older men with low testosterone. Hosp Pract (1995) 46, 47–55 (2018).

29. S. Basaria et al., Adverse events associated with testosterone administration. N Engl J Med 363, 109–122 (2010).

30. S. J. Ohlander, B. Varghese, A. W. Pastuszak, Erythrocytosis Following Testosterone Therapy. Sex Med Rev 6, 77–85 (2018).

31. W. de Ronde, F. H. de Jong, Aromatase inhibitors in men: effects and therapeutic options. Reprod Biol Endocrinol 9, 93 (2011).

32. A. Sansone, F. Romanelli, M. Sansone, A. Lenzi, L. Di Luigi, Gynecomastia and hormones. Endocrine 55, 37–44 (2017).

33. T. Chen et al., Different levels of estradiol are correlated with sexual dysfunction in adult men. Sci Rep 10, 12660 (2020).

34. H. S. Narula, H. E. Carlson, Gynaecomastia--pathophysiology, diagnosis and treatment. Nat Rev Endocrinol 10, 684–698 (2014).

35. M. Schulster, A. M. Bernie, R. Ramasamy, The role of estradiol in male reproductive function. Asian J Androl 18, 435–440 (2016).

36. L. Prokai et al., The prodrug DHED selectively delivers 17beta-estradiol to the brain for treating estrogen-responsive disorders. Sci Transl Med 7, 297ra113 (2015).

37. Lorsch, Z. S. et al. Estrogen receptor α drives pro-resilient transcription in mouse models of depression. Nat. Commun. 9, 1116 (2018).

38. C. E. Terrillion, T. C. Francis, A. C. Puche, M. K. Lobo, T. D. Gould, Decreased Nucleus Accumbens Expression of Psychiatric Disorder Risk Gene Cacna1c Promotes Susceptibility to Social Stress. Int J Neuropsychopharmacol 20, 428–433 (2017).

39. P. Zanos et al., NMDAR inhibition-independent antidepressant actions of ketamine metabolites. Nature 533, 481–486 (2016).

40. P. Zanos et al., Sex-dependent modulation of age-related cognitive decline by the L-type calcium channel gene Cacna1c (Cav 1.2). Eur J Neurosci 42, 2499–2507 (2015).

41. A. Can et al., The mouse forced swim test. J Vis Exp, e3638 (2012).

42. E. Martianova, S. Aronson, C. D. Proulx, Multi-Fiber Photometry to Record Neural Activity in Freely-Moving Animals. J Vis Exp, (2019).

43. Z. M. Zhang, S. Chen, Y. Z. Liang, Baseline correction using adaptive iteratively reweighted penalized least squares. Analyst 135, 1138–1146 (2010).

